# Bilayer-bound structure of SARS-CoV-2 Envelope protein in complex with a host PDZ domain

**DOI:** 10.64898/2026.10.06.756756

**Authors:** Abel Cherian Varkey, Evgeny Nimerovsky, Kai Xue, Claudia Schwiegk, Karin Giller, Stefan Becker, Loren B. Andreas

**Affiliations:** Max-Planck Institute for Multidisciplinary Sciences, Am Fassberg 11, Goettingen, 37077 Germany; Centre of High Field NMR Spectroscopy and Imaging, Nanyang Technological University, 21 Nanyang Link, 637371 Singapore; School of Physical and Mathematical Science, Nanyang Technological University, 21 Nanyang Link, 637371 Singapore

## Abstract

Small multifunctional viral membrane proteins called viroporins are attractive drug targets but are difficult to structurally characterize in a lipid-bilayer environment. We have successfully reconstituted the full-length Envelope (E) protein from SARS-CoV-2 in a lipid-bilayer, co-assembled with a native host factor that binds to E, the PDZ domain 2 of ZO1. We have leveraged the enhancement afforded by dynamic nuclear polarization NMR to determine that the ensemble-averaged oligomer state of E is a pentamer. Complexing the protein with a host PDZ domain did not impact the oligomer state, but resulted in structuring of the last four residues of E, and immobilization of the PDZ domain near the lipid surface. Combined with secondary structure information derived from the chemical shifts, these data were used to model an E protein pentamer together with the PDZ protein dimer. Inclusion of a relevant host factor, as well as a complex lipid bilayer that mimics conditions in the ER-Golgi intermediate compartment, approaches native conditions, and addresses the ongoing controversy regarding the functional oligomer state in an infected cell.

## Introduction

An open reading frame (ORF) towards the 3’-terminus of the single-stranded positive-sense RNA of SARS-CoV-2 encodes its smallest structural protein, the Envelope (E) protein^1^. E is a 75-residue multifunctional membrane protein that is incorporated in low numbers in the mature virus particle. The protein is primarily functional in the infected cell, localizing to the membrane of the endoplasmic reticulum-Golgi intermediate compartment (ERGIC).^2^ E has three structural domains, residues 1-8 representing the luminal N-terminal domain (NTD), residues 9-37 representing the transmembrane domain (TMD), and residues 38-75 representing the cytoplasmic domain (CTD).^3,4^

Within the ERGIC membrane, E is hypothesized to predominantly conduct cations via the TMD with disputable selectivity. Various studies have proposed monovalent (Na^+^, K^+^ and Cs^+^) and divalent cation (Ca^2+^ and Mg^2+^) conduction, as well as Cl^-^ ion conduction depending on the lipid composition and pH.^5–11^ Evidence suggests that E also plays a role in budding by inducing membrane curvature.^12,13^ Coupled with its role in membrane curvature, E is reported to interact with another viral structural protein, the Membrane (M) protein, enabling the formation of virus-like particles (VLPs).^14^

Besides the above-mentioned functions, E participates in protein-protein interaction (PPI) with host proteins through a PDZ domain binding motif (PBM) consisting of the four C-terminal residues.^15^ The importance of the PBM functionality to the virus was elucidated by passaging PBM-deficient mutants of SARS-CoV-1 that gave rise to either reversions or compensatory mutations after several passages of the virus.^16^

PDZ domains, named after the proteins PSD-95, DLG1, and ZO-1, span all kingdoms of life and are therefore common targets for viral interaction to modulate cellular processes as part of the viral infection and lifecycle.^17^ PDZ domains function as PPI modules in proteins that are a part of signal transduction complexes important for cell cycle, cellular communication, cell-cell junction organization and maintenance of cell polarity as well as other roles.^18^ They are 90-100 residues long with a conserved domain architecture. The PBM of an interacting partner is usually the C-terminal four residues, which forms a β-strand that binds in the groove formed by β-strand B and α-helix B of the PDZ domain, although internal PBMs have also been reported.^19^

The PBM of E, –^72^DLLV^75^ is a class-II PBM which is defined by the motif -X-Φ-X-Φ_COOH_, where X represents any amino acid and Φ represents hydrophobic residues.^20^ High throughput experiments have revealed several PDZ domains that interact with E^21–23^ as is characteristic of PBMs.^24^ Of these PDZ domains, the PDZ domain 2 (PDZ2) of the Zonula Occludens 1 (ZO-1), also known as Tight Junction Protein 1 (TJP1), showed the highest affinity towards a C-terminally derived peptide of E.^21,22,25^ We refer to this domain as ZO1-PDZ2 in the remaining text.

ZO-1 is a scaffolding protein present near the inner leaflet of the plasma membrane (PM) as part of the Tight Junction (TJ) complex. The TJ complex is responsible for sealing the gap between epithelial and endothelial cells and regulating paracellular transport.^26^ E, through its interaction with ZO1-PDZ2 is hypothesized to mis-localize ZO-1 from the PM to the ERGIC membrane. Mis-localized ZO-1 in turn is conjectured to destabilize the TJ complex, leading to vascular leakage, and consequently aid the spread of the virus inside the lungs. This mechanism, along with ion conduction, is invoked to explain aspects of the acute respiratory distress syndrome (ARDS) observed in severe cases of COVID-19, the respiratory disease caused by SARS-CoV-2^7^ and is being explored as a potentially druggable interaction.^27^

Structurally, ZO1-PDZ2 forms a domain-swapped homodimer with two binding pockets for PBMs.^28^ Due to its ability to dimerize, it is involved in crosslinking different proteins in the TJ complex. ZO1-PDZ2 displays similar binding modes with native^29^ and viral PBMs.^27^

While C-terminally derived peptides of E have been structurally characterized in complex with ZO1-PDZ2 as well as other PDZ domain proteins,^25,30^ information about the rest of E has focused on various truncated constructs in the absence of PDZ domain proteins. To date, five experimentally determined structures of E have been deposited to the Protein Data Bank (PDB). Of these five, two structures are of E from SARS-CoV-1, which has a 95% sequence similarity to SARS-CoV-2 E, and is identical to SARS-CoV-2 E in the NTD and TM domains. These two structures were determined using solution NMR from a truncated E construct spanning residues 8-65 (ETR), one in SDS micelles^31^ and the other in LMPG micelles.^32^ The remaining three constructs are TM constructs of E determined using solid-state NMR. Of the three, one spans residues 12-37 (ETM_12-37_) reconstituted in a POPC/POPG bilayer.^33^ The remaining two are reconstituted in an ERGIC membrane-mimetic composed of five lipids, which we refer to as ERGIC-lipids (see Methods), and span residues 8-38 (ETM_8-38_) representing closed^34^ and open^35^ conformations prepared near neutral pH, or in the presence of calcium and low pH, respectively. Thus, atomic resolution studies of E have categorically delinked the TM domain responsible for ion conduction and the C-terminus responsible for PDZ binding.

Another unresolved critical facet of E pertains to the oligomeric state of the protein, which has particular significance for ion conduction since oligomer interfaces define the TM structure and therefore the potential pathways for ions. The oligomeric state of E is modulated by the membrane environment and the length of the construct used. LMPG micellar ETR was modelled as a pentamer based on earlier experimental results. A construct of E containing the N-terminal residues, spanning residues 1-41 (ENTM), reconstituted in DMPC/DMPG,^36^ along with ETM_12-37_ reconstituted in POPC/POPG, were reported to be dimeric.^33^ ETM_8-38_ was prone to clustering in ERGIC-lipids, while the removal of phosphatidylcholine (PI) and cholesterol lipids from the ERGIC-lipids mixture suppressed clustering and showed a pentameric state.^37^ Full-length E (EFL) was reported to be a dimer in various detergents using native mass spectrometry while the shoulder in a mass photometry profile of EFL in nanodisc as well as sedimentation velocity experiments of EFL in C14-betaine detergent were interpreted as pentamers.^38–40^ Recently, size exclusion chromatography multi-angle light scattering (SEC-MALS) was used to show that EFL reconstituted in POPC/POPE/POPS nanodiscs formed by MSP1D1ΔH5 is pentameric.^41^

In this report, we measure the oligomeric state of EFL in complex with ZO1-PDZ2 as well as elucidate details about secondary structure and dynamics of EFL and ZO1-PDZ2 within the complex. This information is combined to generate a pentamer model of the multifunctional E protein in complex with a hijacked host PDZ domain.

## Results

### MAS NMR reveals a specific interaction between the PBM of E and ZO1-PDZ2 dimer

We first expressed and reconstituted ^15^N-labelled EFL in a wide array of lipid compositions (Supplementary Figure 1). The CP based ^1^H-^15^N correlation spectra showed overlapping broad resonances with minor differences between the different samples. Given only subtle differences between spectra in the various lipid compositions, we chose the biologically relevant ERGIC-mimetic mixture of five lipids for further experiments.^34,42^

Both SDS-PAGE gels and NMR spectra confirm successful reconstitution of PDZ binding for EFL in lipid bilayers (Figure 1). Figure 1A shows details of the interaction between ZO1-PDZ2 and a twelve-residue C-terminally derived peptide of E from a 1.72 Å crystal structure.^27^ The PBM of E forms an anti-parallel β-strand aligned next to and interacting with the βB strand of ZO1-PDZ2 through backbone mediated hydrogen bonds involving residues V70, L73 and V75. The terminal -COOH moiety of this peptide forms additional hydrogen bonds with the ‘GLGF’-motif present in the loop region between βA and βB strands of ZO1-PDZ2. Figure 1B confirms the interaction via SDS-PAGE, and Figure 1C-E shows NMR spectra that identify the interacting residues of E. Insertion of EFL into the bilayer membrane was confirmed by obtaining ^1^H-^1^H NOESY cross-peaks between E and mobile lipid molecules (Supplementary Figure 2C).^43^ Additionally, transmission electron microscopy (TEM) images of the reconstituted proteoliposomes showed speckles that are interpreted as protein complexes. (Supplementary Figure 2D).

**Figure 1.**
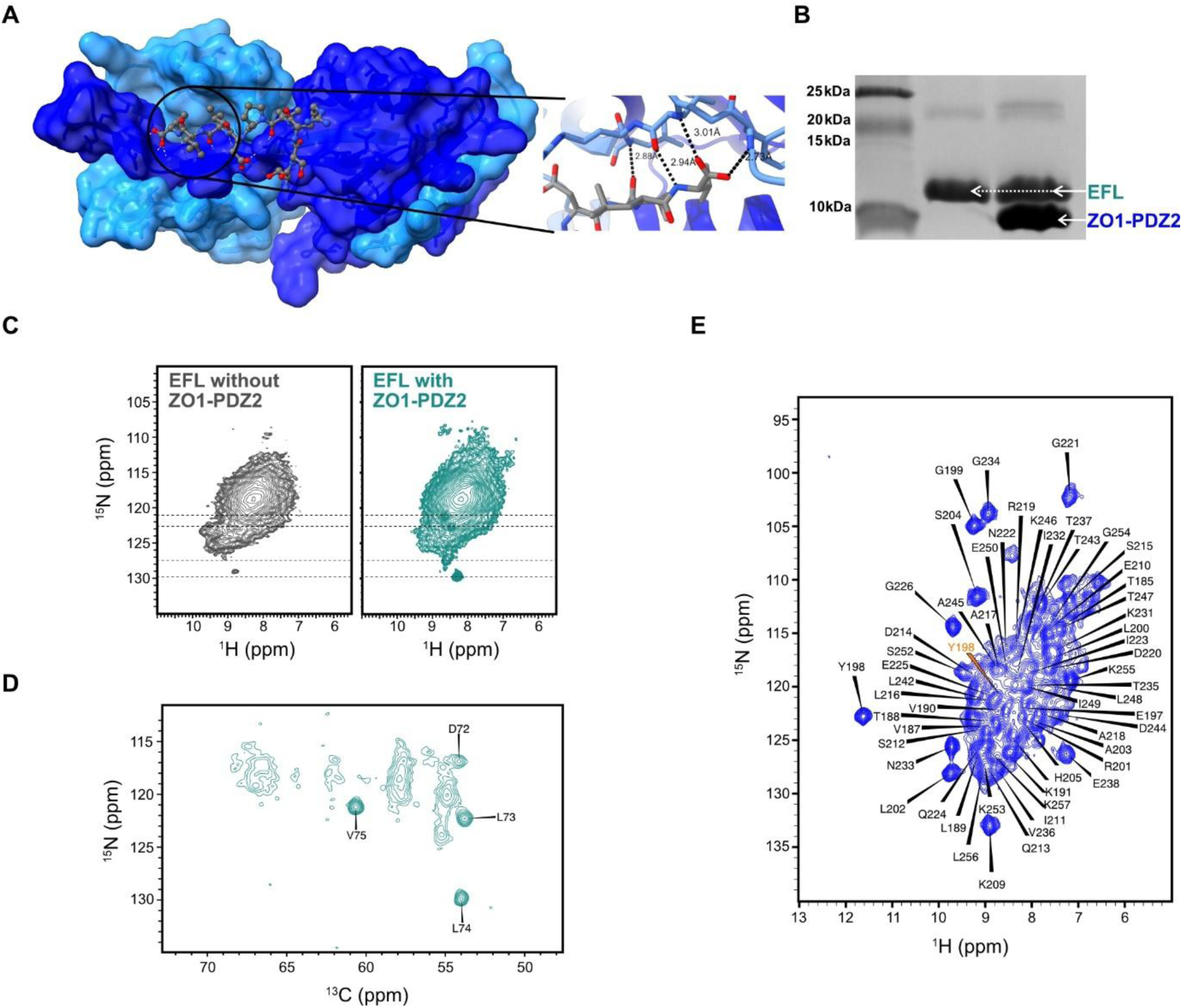
Spectroscopic signatures of EFL and ZO1-PDZ2 interaction via the PBM. **(A)** Crystal structure of ZO1-PDZ2 in complex with a twelve-residue C-terminal peptide of E (pdb_00009HM2). The two chains of ZO1-PDZ2 are shown as surface models, colored light and dark blue. The twelve-residue peptide is shown as a ball and stick model in grey. The expansion displays the PBM interaction highlighting distances between carbonyl O and amide N present between hydrogen-bonded residues. **(B)** SDS-PAGE showing the reconstituted EFL without and with ZO1-PDZ2. **(C)** The ^1^H-^15^N correlation spectra of EFL without and with ZO1-PDZ2 bound. The dotted lines indicate peaks that appear only in the presence of ZO1-PDZ2. (**D)** The ^13^C-^15^N projection from an (H)CANH spectrum showing the four sharp peaks corresponding to the PBM of EFL acquired at 1200 MHz at 55 kHz MAS. **(E)** The ^1^H-^15^N correlation spectrum of ZO1-PDZ2 with assignments. ZO1-PDZ2 is in complex with EFL reconstituted in ERGIC-lipid bilayer. Spectrum acquired at 800 MHz and 55 kHz MAS. The peak position of Y198 (BMRB 16871) is displayed in orange.

Strikingly, four sharp peaks are observed in the ^1^H-^15^N correlation spectra of ^2^H, ^13^C, ^15^N-labelled EFL reconstituted with ZO1-PDZ2. These peaks were absent from EFL reconstituted without ZO1-PDZ2 (Figure 1C). Beginning from the downfield shifted terminal –COOH peak at 180.3 ppm, the backbone resonances of ^72^DLLV^75^ were readily assigned (Figure 1C, Supplementary Table 1) using a suite of assignment spectra (see Methods). The chemical shifts of the terminal three residues lie in the expected β-strand chemical shift range while D72 shows more intermediate coil-like chemical shifts.

To characterize ZO1-PDZ2 in complex with EFL in a bilayer context, we reconstituted ^2^H,^13^C,^15^N-labelled ZO1-PDZ2 with unlabeled EFL. The CP based ^1^H-^15^N correlation spectrum of ZO1-PDZ2 showed many well-resolved peaks (Figure 1E). The result was surprising since soluble proteins such as ZO1-PDZ2 have short molecular tumbling correlation times when free in solution, which interferes with CP-based polarization transfer. These data indicate that ZO1-PDZ2 is well folded and relatively immobile on the bilayer surface through interaction with E PBM with a residence time on the order of milliseconds or longer. When the ZO1-PDZ2 was reconstituted in the same lipid composition without EFL, we obtained an unresolved weak spectrum, possibly from precipitated or partially folded protein (Supplementary Figure 3C).

Approximately 60% of the backbone residues of ZO1-PDZ2 were successfully assigned from a set of seven spectra (Supplementary Figure 4).^44^ The comparison of the solution assignments of ZO1-PDZ2 in the absence of E PBM (BMRB entry 16871) revealed differences in several peaks close to the binding site compared to our assignments of bound ZO1-PDZ2 as shown by chemical shift perturbations (CSP) (Supplementary Figure 5). Some of the differences may arise due to differences in pH and construct used in the two studies. The largest CSP observed for Y198 corroborates the hydrogen-bonding between V75 of E and Y198 of ZO1-PDZ2 inferred from the crystal structure (Figure 1A). Given that ZO1-PDZ2 is a well-folded domain, we do not expect large overall structural differences compared to the protein in crystalline form and did not pursue further structural characterization of the protein.

Consistent intensities across the peaks of ZO1-PDZ2 involved in binding E PBM compared to the rest of the protein suggest that the majority of the detected ZO1-PDZ2 has both sites occupied by the E PBM. This implies that either the population of ZO1-PDZ2 dimers with singly occupied binding sites is low, or the singly-occupied ZO1-PDZ2 proteins are dynamic on a timescale invisible to cross polarization (CP)-based NMR. Outside the PBM region, the solution and MAS-NMR assignments of the homodimeric ZO1-PDZ2 are consistent.

We also recorded static ^31^P spectra to probe changes to the lipid bilayer in the presence and absence of ZO1-PDZ2, but no significant changes were seen (Supplementary Figures 2A and 2B).

**Figure 2.**
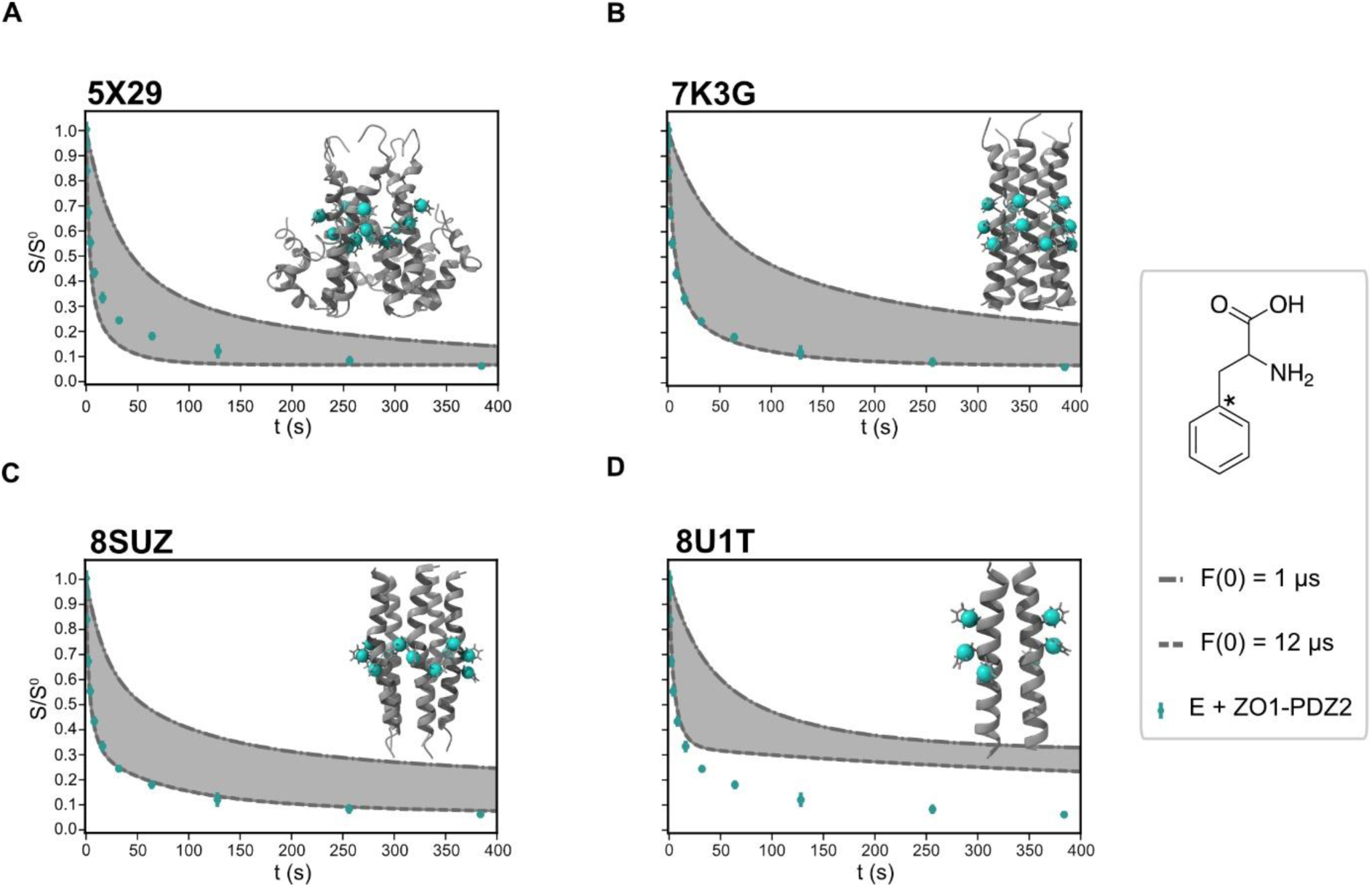
Experimental ^13^C-CODEX of EFL with ZO1-PDZ2 and CODEX simulations. Structures of E highlight the labelled Cγ atoms of the phenylalanine residues as cyan spheres. For each panel, the background-corrected CODEX data acquired for EFL in complex with ZO1-PDZ2 is displayed as cyan points. The grey shaded region indicates the area bounded by CODEX curves simulated for the given model with 1 μs and 12 μs, which corresponds to minimum and maximum reported F(0) values, respectively. The legend for all panels is displayed on the right. **(A)** ETR from SARS-CoV-1 in LMPG micelles, **(B)** the closed ETM_8-38_ model from SARS-CoV-2 in ERGIC-lipids, **(C)** the open ETM_8-38_ model from SARS-CoV-2 in ERGIC-lipids, and **(D)** the dimeric ETM_12-37_ from SARS-CoV-2 in POPC/POPG lipids. CODEX data were acquired at 600 MHz at 8 kHz MAS and 92 K.

#### Oligomer state of E

After the successful reconstitution of EFL and ZO1-PDZ2 in ERGIC-lipids, we sought to assess the oligomeric state of EFL, in the absence and presence of its host binding partner, ZO1-PDZ2. The MAS-NMR technique Centerband-Only Detection of EXchange (CODEX) can be used in conjunction with tailored labeling to determine oligomer state.^37,45–48^ Briefly, CODEX measures the dephasing of chemical shift anisotropy (CSA) encoded NMR signal due to a change in CSA tensor orientation that occurs during the mixing time.^49,50^ For investigation of oligomers, the reorientation occurs due to spin diffusion within a small set of spins corresponding to labelled sites of the oligomer. The equilibrium value achieved after a sufficiently long mixing period is equivalent to the reciprocal of the number of neighboring spins. Usually, the number of spins per monomer is restricted to one via selective labelling and subsequently the count of spins obtained by CODEX is equal to the oligomer number. Otherwise, the oligomer number is obtained by simply scaling by the number of spins per monomer.

Since ^19^F is naturally absent in biological samples and has a large gyromagnetic ratio (γ), ^19^F has been the preferred spectroscopic probe for oligomer determination of membrane proteins using CODEX. Despite its advantages, introducing ^19^F-labelled amino acids into recombinant proteins with a high labelling efficiency is challenging.^51^ We, therefore, carried out ^13^C-based CODEX with DNP to compensate for the low gyromagnetic ratio of ^13^C and to enhance the starting polarization.^52^

We selected phenylalanine residues for labelling, since the large CSA of aromatic moieties facilitates CODEX measurements, and since there is minimal overlap with lipid signals. The sequence of E contains five phenylalanine residues, three of which are in the transmembrane region expected to oligomerize. We therefore generated an F4Y/F56V double mutant of E such that phenylalanine residues appear thrice in the sequence and are restricted to the TM domain of the protein. F4Y was chosen to retain aromaticity and to have similar space filling while F56V was chosen based on the presence of V56 in the sequence of SARS-CoV-1 E. ^13^C-γ-phenylalanine labelling was introduced for its favorable CSA and T_1_ as shown earlier with GB1.^53^

CODEX measurements were carried out on three samples: EFL in complex with ZO1-PDZ2 at a lipid to protein molar ratio (LPR) 20 (Figure 2), and EFL without ZO1-PDZ2 at two lipid concentrations of LPR 20 and 30 (Supplementary Figure 6). As mentioned earlier, since there are three phenylalanine residues per F4Y/F56V EFL monomer, the S/S_0_ equilibrium value reached by the CODEX curve at long times approaches a reciprocal of three times the oligomer number, where S and S_0_ correspond to intensities of the dephased and reference curves, respectively. The three CODEX curves reach equilibrium values consistent with a pentameric protein and a small degree of clustering of pentamers. In keeping with this interpretation, EFL without ZO1-PDZ2 at an LPR of 20, dephased more in the CODEX measurements as compared to LPR of 30, while EFL with ZO1-PDZ2 at LPR 20 is in between the two. These subtle differences are consistent with screening of pentamer-pentamer contacts in the membrane by both lipids and ZO1-PDZ2. The results thus suggest that EFL forms a pentamer both with and without ZO1-PDZ2 in the complex.

The CODEX decay curves were simulated for the four PDB structures shown in Figure 2 using the well-established rate matrix approach.^46^ The rate of magnetization exchange between the labelled phenylalanine Cγ sites depends on the dipolar coupling between the sites, dictated by the protein structure, as well as an additional F(0) term. F(0) represents the probability that single quantum transitions for two spins occur at the same frequency. In practice, F(0) is measured for model compounds of known structure rather than being calculated *ab initio*. The measured value is then applied for distance measurement in samples of unknown structure. The grey shaded regions in the plots in Figure 3 are bounded by simulated curves with F(0) between 1 and 12 μs which are approximately the highest and lowest values reported in literature for comparable ^13^C spins.^54^ In the case of triply-labeled EFL, either six sites in a dimer or fifteen sites in a pentamer are involved in magnetization exchange. The equilibrium value for the pentamer structures of Figure 2A-C is therefore about 6.7 percent, in agreement with the data (6.2±0.5%) while the simulations of Figure 2D reach only about 17 percent.

**Figure 3.**
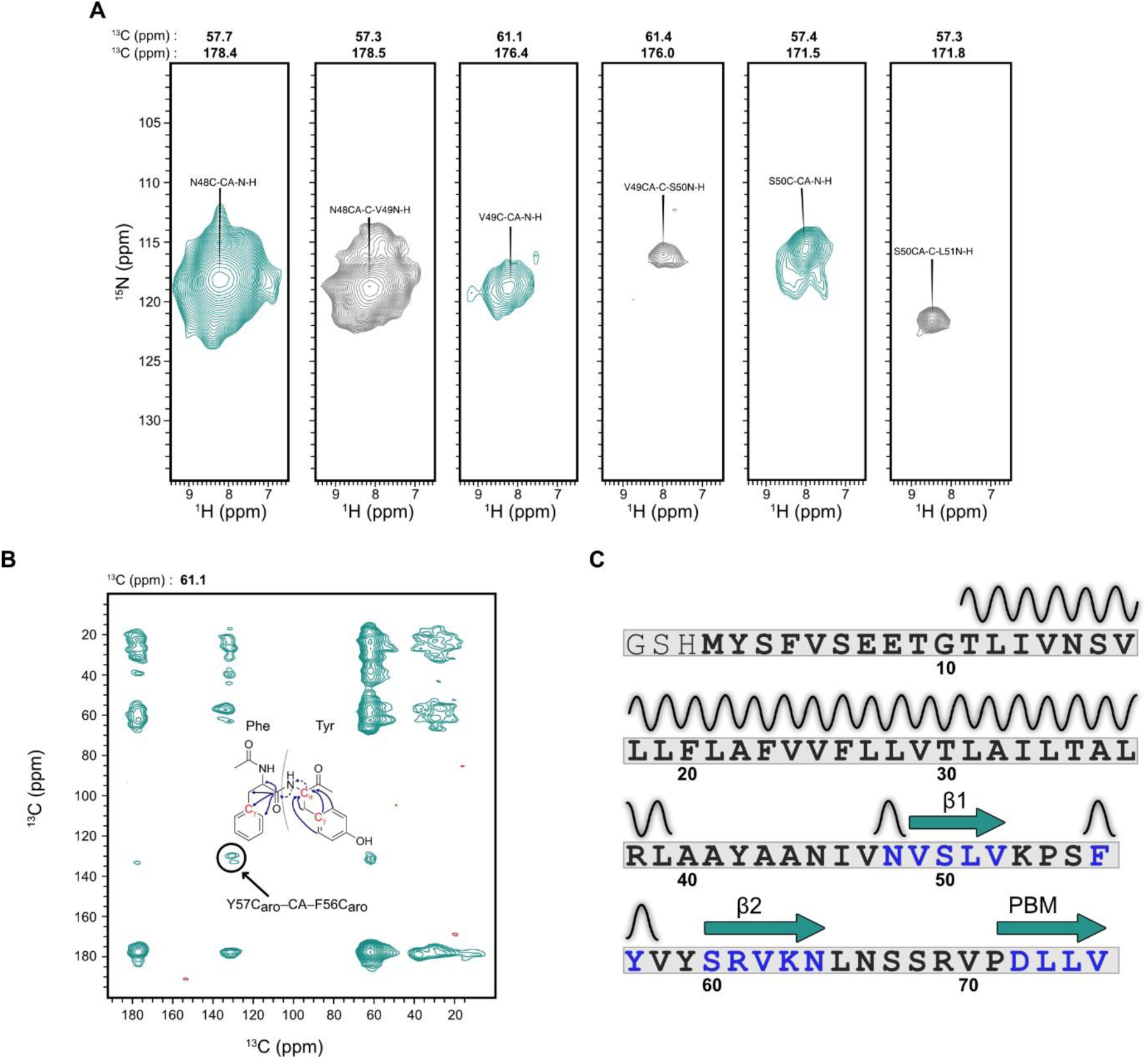
Assignment of EFL with ZO1-PDZ2 based on the 4D spectra and DNP. **(A)** Strips from (H)COCANH are displayed showing the assigned resonances of the CTD. N-H plane is shown with the Cα and CO resonance frequencies mentioned above the spectra. **(B)** (H)CXCA(N)(CO)CX magnetization pathway and the corresponding spectrum obtained with DNP enhancement is shown. The F56-Y57 cross-peak is shown. The spectrum was acquired at 600 MHz at 12.5 kHz MAS under DNP conditions at 92 K. **(C)** The sequence of EFL is displayed with assigned residues in blue, and helical (wavy line) or beta sheet (green arrows) secondary structures identified based on chemical shifts. The initial three residues (GSH) are cloning artefacts and are not included in the sequence numbering.

Based on the distribution of CODEX data points that lie within this shaded region, the ETR structure in LMPG micelles from SARS-CoV-1 shows the most feasible arrangement of phenylalanine residues. This arrangement has the shortest interchain phenylalanine distance. The CODEX data of EFL at an LPR of 30 also fit within the bounds of the open ETM_8-38_ CODEX simulation (Supplementary Table 2). In summary, the CODEX data support a pentameric arrangement with relatively close phenylalanine contacts compared with the set of reported PDB structures of E.

#### Assignments and secondary structure of E

Although the PBM resonances of EFL showed ∼0.5 ppm ^13^C and ∼1 ppm ^15^N linewidths under the tremendous resolving power of the 1.2 GHz magnet, the rest of the protein resonances displayed ∼1.2 ppm ^13^C and ∼2 ppm ^15^N linewidths, but were difficult to measure due to significant peak overlap. Poor spectral resolution of EFL in complex with ZO1-PDZ2 limited access to detailed structural information across the whole protein. Even with the combined benefits of ^1^H-detected experiments at 55 kHz MAS on a perdeuterated sample, it is challenging to obtain assignments of EFL using standard 3D assignment sequences.

Nevertheless, buoyed by recent developments in sensitivity enhanced homonuclear and heteronuclear polarization transfer elements, we recorded 4D assignment spectra using (H)CACONH and (H)COCANH pulse sequences.^55,56^ The 4D datasets enabled the assignment of two stretches of residues in the CTD of EFL, –^48^NVSLV^52^– and –^58^YSRVKN^63^– (Figure 3A, Supplementary Table 1). We also used the S3 pulse sequence,^57^ implemented as (H)CXCA(N)(CO)CX in conjunction with DNP to obtain the Cα chemical shift of Y^57^ (Figure 4B). We also recorded an (H)COCANH spectrum of EFL without ZO1-PDZ2 reconstituted in ERGIC-lipids and observed the well-isolated S50 peak (Supplementary Figure 7).

**Figure 4.**
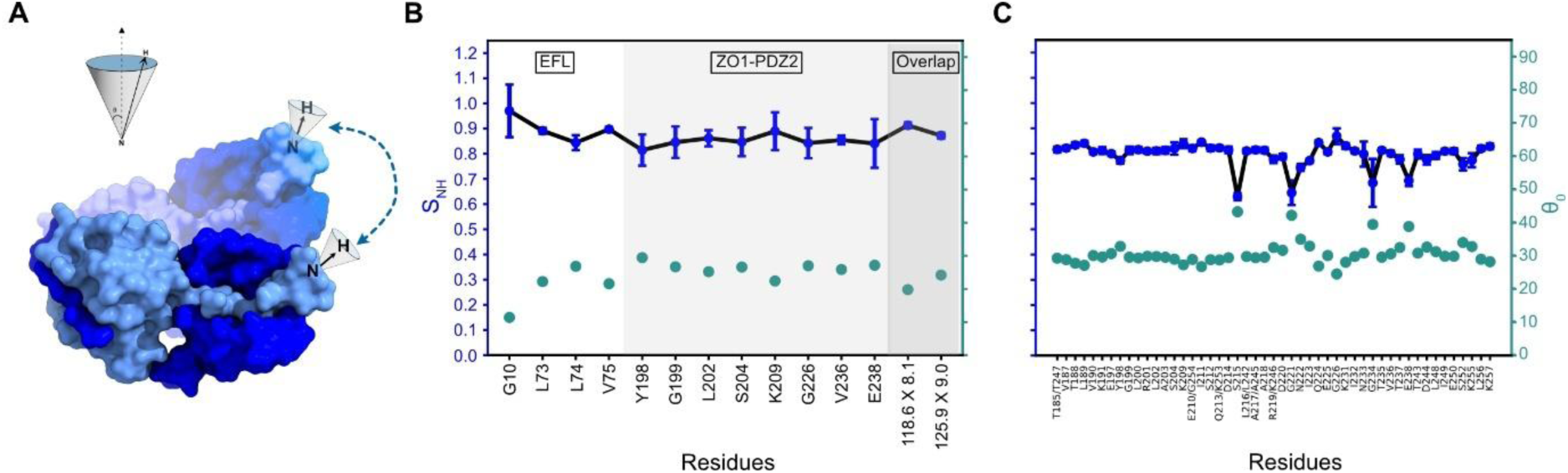
Sub-microsecond dynamics from ^1^H-^15^N single bond dipolar coupling measurements. **(A)** A protein (pdb_00009HM2) is displayed showing rocking motion about an arbitrary axis. The NH bond vector shown on the protein changes orientation due to the overall protein motion as well as local motion. The inset shows diffusion in a cone as an example of one such motional model for the rocking motion with the amplitude of the motion captured by the cone half angle. **(B)** Measured ^1^H-^15^N dipolar coupling values for ^2^H,^13^C,^15^N-labelled EFL in complex with ^15^N-lablled ZO1-PDZ2 converted to S_NH_ and displayed in blue, as a function of residues of EFL, ZO1-PDZ2 (grey shading) and for overlapped peaks (dark grey shading). If the motional model is assumed to be diffusion in a cone, the S_NH_ values can be converted to cone half angles that are displayed in green. **(C)** Measured ^1^H-^15^N dipolar coupling value for ^2^H,^13^C,^15^N-labelled ZO1-PDZ2 in complex with unlabeled EFL displayed as S_NH_ and cone half-angles. Data were acquired at 800 MHz at 55 kHz MAS.

These chemical shift values provide information about the secondary structure in the CTD, as summarized in Figure 4C. TALOS-N was used to predict the torsion angles based on the chemical shifts.^58^ The torsion angles of N48 fit better to a helical secondary structure that likely marks the end of the amphipathic helix seen in the micellar ETR structure.^32^ Subsequently, the residues transition to an extended sheet conformation from V49 until V52. A second stretch of extended residues from S60-N44 is separated from the first by five unassigned residues and the unique aromatic-aromatic pair, –^56^FY^57^–, with more helical chemical shifts. At 62 ppm, the Cα chemical shift of Y57 clearly shows a non-extended secondary structure. These assignment data provide detailed insights into the secondary structure of the E protein CTD at different points along the sequence, which can then be compared with different proposed structures of the protein, data on truncated E protein in different environments, as well as proposed structural models (see discussion).

#### Dynamics of E and ZO1-PDZ2

The observation of ZO1-PDZ2 in Figure 1E indicates that the pentameric EFL quenched rotational diffusion of ZO1-PDZ2 via interaction with the PBM residues. To quantify the fast dynamics in the complex, we measured ^1^H–^15^N single-bond dipolar coupling strengths using the pMODERN recoupling pulse sequence.^59^ When bond angular motions occur on timescales shorter than ∼100 ns,^60^ the measured dipolar coupling strength scales inversely with the amplitude of the motion (Figure 4A).

The pMODERN results indicate that both EFL and ZO1-PDZ2 are similarly rigid, with an order parameter S_NH_ of about 0.85. The order parameter, S_NH_, is normalized against the rigid limit dipolar coupling such that 1 represents perfectly rigid and 0 represents complete averaging of the coupling. The parameter S_NH_ depends on the amplitude of motion and the motional model. If we assume that the motion of the dipolar vector can be represented as a diffusion in a cone, then S_NH_ can be converted to the cone half-angle as shown in Figures 4B and 4C.^61^ The low-amplitude total motion of the NH bond vector detected by dipolar coupling measurements largely correspond to cone half-angles between 20° and 30°.

We measured the ^1^H-^15^N single-bond dipolar couplings from two samples. In the first sample, both EFL and ZO1-PDZ2 are ^15^N-labelled. Both isolated and overlapped peak positions were analyzed from this sample. The isolated peaks indicated that both proteins in the complex are relatively rigid. Since the overlapped peaks are a convolution of two different proteins, as well as several sites within E, they represent an average behavior across the complex, which again indicated close to rigid values for S_NH_. The second sample had ^15^N-labelling only on ZO1-PDZ2 that allowed us to obtain site specific dipolar coupling strengths from the better-resolved ZO1-PDZ2 spectrum. The higher mobility of loop residues S215, G221, G234 and E238 indicates higher local flexibility outside regions of regular secondary structure. Both samples indicate that ZO1-PDZ2 is tethered adjacent to the bilayer with heavily restricted motion.

## Discussion

Research on the structure of E until now has separated the PBM functionality from the rest of the protein. In this report, we bridge this gap by characterizing the structure of EFL, bound to a host PDZ domain, reconstituted in bilayer lipids matching the ERGIC-membrane composition.

### Oligomer state of E

The measured oligomer state of E depends on the length of the construct, reconstitution environment, protein concentration and the technique used. Although a range of oligomeric states have been observed, especially in SDS-PAGE gels, experiments from bilayer environments or reconstituted in milder detergents indicated either a dimer or a pentamer.^11,33,36,37,62,63^

Dimers have been reported in micellar detergent preparations using native mass spectrometry and analytical ultracentrifugation.^11,38^ Dimers of E in liposomal preparations were shown for the ETM_12-37_ construct at an LPR of 25 in POPC/POPG bilayer^33^ and ENTM at LPR 30 in DMPC/DMPG.^36^ Three arguments have been put forth to support dimeric E: the interaction of E with the dimeric M protein would be facilitated by a corresponding dimer,^36^ the low level of E in a mature virus particle might promote dimer over pentamer,^64^ and lastly, that E exchanges between a storage dimer and functional pentamer pool in the ERGIC membrane.^65^ Despite these reasons, to selectively conduct ions, E must be capable of forming a pore in the membrane which is justified by the pentameric organization. Dimeric E can perform ion conduction only through membrane disruption, but this would be at the cost of directionality and selectivity of ion conduction, not excluding the distinction between cations and anions. Our CODEX results rule out a predominantly dimeric population and are instead consistent with pentameric E in the ERGIC bilayer.

The presence of ZO1-PDZ2 does not alter this oligomer state. The dimeric ZO1-PDZ2, surprisingly, does not correspond to a submultiple of pentameric E, raising the possibility that a fraction of the pool of ZO1 protein is bound to E with only one monomer of the dimer. A second possibility that cannot be ruled out from our data is the presence of both dimers and oligomers of E larger than pentamers. However, such a possibility would diverge from the preponderance of literature indicating either dimeric or pentameric forms of the protein.

As inferred from the CODEX experiments, the dilution of EFL with additional lipids does decrease oligomer clustering but does not significantly alter the ensemble-averaged oligomer number. Negative-stain TEM images of our liposome preparations indicate a preference for the protein complex to cluster in regions of the liposome rather than a homogenous distribution of particles (Supplementary figure 2D). Large regions of the liposomal surface are seemingly free of proteins in the images. This would suggest that even higher dilutions would be required to shift the equilibrium away from pentamers, further corroborating the conclusion that a pentameric E is the relevant form in an infected cell. The fact that dimeric PDZ can interact with two molecules of E belonging to separate oligomers can also be expected to increase the local concentration of E that is further discussed later.

Inversely, a higher local concentration of E naturally strengthens the PDZ interaction through multivalent interactions. The techniques used to characterize the affinity between the PBM of E and ZO1-PDZ2 have relied on using C-terminal derived peptides of E that have commonly reported affinities in the micromolar range, although one report estimated nanomolar binding affinity.^21^ These relatively weak affinities are monovalent interactions with respect to E, and suggest that the oligomerization of both proteins may play an important role in strengthening the overall interaction.

### EFL reconstituted in a lipid bilayer

Our experience with Fos-Choline-16 solubilized EFL showed that the PDZ interaction is screened by the presence of the detergent, in contrast with lipid reconstituted EFL which faithfully reproduces the interaction. Monomeric detergent found in micellar conditions is most likely responsible for this screening (Supplementary figure 3A). This explanation fits the hydrophobic character of the binding pocket that interacts with a PBM comprised of at least two hydrophobic residues. More broadly, the TM helices of viroporins have also been shown to be sensitive to the solvating membrane or membrane-mimetic environment, which has important implications for understanding the conduction mechanism.^66,67^

The phenylalanine residues that are part of the TM domain have been proposed to be involved in ion conduction as part of an aromatic network involved in concerted motion during channel opening.^68^ The F(0) values extracted from the CODEX curve is a good indicator of feasible models, even if it does not provide precise high-resolution information due to the contribution from multiple unequal spin-spin distances and orientations. The results we obtained are most consistent with the arrangement of phenylalanine residues reported in a solution NMR structure (pdb_00005X29) in which F26 is found inside the pore. Other pentameric structures deposited in the PDB cannot be excluded but would require a spin diffusion rate that slightly exceeds the fastest rate observed in model systems (Figure 2). It has also been proposed that F20 and F26 residues have different χ_2_ angles in the closed ETM_8-38_ as compared to the deposited PDB structure (pdb_00007K3G), but the same χ_1_ angles as the deposited structure.^68^ Since the position of the labelled Cγ site remains unaffected by a change in the χ_2_ torsion angle, this change would not alter the conclusion that the inward F26 pentamer is the best fit to the CODEX data.

### In between functional motifs

As emphasized earlier, high resolution MAS-NMR structures of the ion conducting TMD have been successfully determined from ETM constructs whereas the PBM structure has been revealed from crystal structures of C-terminally derived peptides of E with various PDZ domains. The ∼30 intervening residues have relatively sparse information with conflicting results obtained from ETR constructs that are missing the last 10 residues. The ETR (SARS-CoV-1) structure from LMPG micelles^32^ suggests a largely helical CTD besides a 6-residue extended region. Meanwhile, secondary structure information gleaned from ETR (SARS-CoV-2) in DMPC/DMPG and ERGIC-lipids suggests three β-strands in the CTD.^40,64^ Our data on EFL in complex with a native host factor, as inferred from the chemical shifts, shows a mixture of α-helical and β-sheet structures in the CTD.

The spectra of EFL reported here do not show a majority of the characteristic peaks associated with β-sheet structure observed in ETR in lipids (referred to as liposomal ETR).^64^ Thus, the absence of the last ten residues in ETR appears to influence the structure in the CTD of E, allowing a β-sheet arrangement not seen in the full-length protein (Supplementary Figure 8). The length of the β1-strand in EFL is shorter than the proposed β1-strand of liposomal ETR. The assignments for the residues prior to N48 could not be obtained since the backbone walk leads to the most overlapping region of the spectra corresponding to helical chemical shifts. This observation fits with the amphipathic helix terminating at V47, as modelled in the micellar ETR (SARS-CoV-1) structure determined from LMPG micelles. This region in the native sequence contains three cysteine residues (C40, C43 and C44) that are anchored to the membrane via palmitoylation.^63^ The proposed β2-strand in liposomal ETR is also contestable based on the result of our (H)CXCANCOCX spectrum which places the helical Y57 right in the middle of a helical segment, similar to the micellar ETR structure. The chemical shifts also indicate a β3-strand in EFL that is similar to liposomal ETR but not observed in micellar ETR. Lastly, although we expected the PDZ binding to influence the structure beyond the immediate vicinity of the PBM, the presence of the same S50 peak with (Figure 3A) and without ZO1-PDZ2 (Supplementary figure 7) suggests that the secondary structure of the CTD (excluding the PBM) is largely unaffected by the interaction.

The mixed results demonstrate that truncated constructs are indeed useful initial reporters for the supra-architecture and atomistic structure of proteins, although the precise perturbations due to truncation of the protein and the membrane mimetic, amongst other factors, are difficult to predict. The importance of including additional domains is more pronounced when allosteric coupling exists between the intra- and extramembrane domains of membrane proteins. Examples of allosteric coupling in viral membrane proteins include the amphipathic helix sequence-dependent structural ordering of the TM helix in M2 (Influenza A),^66^ conformational coupling between the TMD and the antigenic surface presented by the cytoplasmic tail domain of the envelope spike protein (HIV-1),^69^ and as seen in this study for E (SARS-CoV-2), the switch from pentameric organization of the TMD,^37^ to dimer upon addition of the NTD,^36^ and back to pentamer upon the successive addition of the CTD.

### Dynamics in the complex

The observed NH dipolar coupling strengths of the amide backbone atoms suggest a relatively rigid EFL and ZO1-PDZ2 complex. The S values when converted to diffusion in a cone model indicate that the motion occurs with an average total amplitude of ∼30°. Previous measurements with chicken α-spectrin SH3 domain microcrystals, for which similar S values close to 0.9 have been observed suggests that such a significant rocking motion for the complex as a whole is unlikely.^70^ It is obvious that overall rocking motion would degrade the quality of X-ray diffraction data but high resolution structures of the SH3 domain exist.^71^ Furthermore, rocking motion in crystals of ubiquitin have been shown to be on the order of a few degrees occurring on tens of microseconds timescale which would not impact the dipolar coupling measurement.^72^ α-helical proteins like AqpZ and M2 show high order parameters for C^α^-H^α^ and N-H^N^ single bonds respectively indicating rigid backbones.^73,74^ A complex of GB1 with the antibody IgG also showed S values of approximately 0.84 for the NH bond.^75^ Therefore, the order parameters seen for EFL in complex with ZO1-PDZ2 can be interpreted mostly as the result of local bond fluctuations including bond stretching and bending, and in addition, possibly very low amplitude librational modes of larger domains, up to the overall complex.

### CODEX for oligomer determination

The success of ^13^C CODEX for a complex of ∼18×5 kDa reconstituted in lipid bilayers was made possible by the signal enhancement of DNP. It is further facilitated by cryogenic temperatures that quench molecular motions, minimizing longitudinal relaxation. Together, this permitted the extensive CODEX mixing as well as five transfer steps in the (H)CXCA(N)(CO)CX assignment experiment. These experiments are currently infeasible for conventional MAS-NMR without hyperpolarization.

In interpreting the oligomer state of E, we also note the limitations of the CODEX technique. Since CODEX experiments report on the ensemble averaged oligomeric state, a mixture of higher and lower oligomeric states would appear as an average oligomer state. Secondly, since the sites chosen for labelling for CODEX experiments by us, as well as by Hong and coworkers,^35–37^ are in the middle of the TM helix, the measurement would also detect head-to-tail packing interactions. Such an arrangement is not expected, based on the physiologically relevant orientation of E in the membrane. Furthermore, the additional bulk from the CTD of EFL as well as the bound ZO1-PDZ2 potentially sterically occludes the close-approach of nearby E oligomers, reducing the risk that CODEX detects such non-specific interaction. Nevertheless, head-to-tail packing interactions being detected by CODEX cannot be completely ruled out.

### Modeling of the EFL and ZO1-PDZ2 complex

We have established that EFL forms pentameric assemblies that form complexes with dimeric ZO1-PDZ2 domains. Using this information along with other structural details gleaned from the experiments and previously reported data, we modeled the structure of the complex. Five copies of EFL and one dimer unit of ZO1-PDZ2 were included. Initially we attempted to generate models where both ZO1-PDZ2 binding sites were occupied, either by two neighboring chains of E or two alternate chains of E, but this led to physically unrealistic models. Based on our observation that the cytoplasmic domain could potentially be preorganized in a β-stranded conformation prior to ZO1-PDZ2 binding, all monomers in the complex were modelled with similar restraints except the last eight residues of the bound monomer as explained next. To simplify the structure calculation in CYANA as well as for better visualization, the first 7 residues of E were excluded from the modelling. Similarly, the last 8 residues of the four unbound monomers of E were also excluded.

The TMD of E was modelled exactly as the TMD of LMPG micellar ETR based on the match between the phenylalanine arrangement and the CODEX data. The proposed amphipathic helix from L38 to N48 that is also seen in the LMPG micellar ETR model has been included in our model in line with EPR data that predicts membrane insertion for this stretch of residues.^76^ The residues from V49 to L51 as well as from S60 to N64 were included as β-strands 1 and 2, respectively, based on chemical shift information (Figure 3). Since β-strands rarely exist in isolation, we explored three possibilities for β-sheet formation. The first possibility involved the β-strands folding back to form an intra-monomer anti-parallel arrangement (Figure 9A), while the two other possibilities involved parallel β-sheet formed by the β1 of one monomer interacting with β2 of a neighboring chain in either direction (clockwise and anticlockwise, Figures 9B and 9C). Residues S55 to V58 were modelled as a single helical turn based on the results of (H)CXCA(N)(CO)CX experiment (Figure 3B). Finally, the PBM interaction with ZO1-PDZ2 as well as the structure of ZO1-PDZ2 dimer was obtained from the crystal structure.

**Figure 9.**
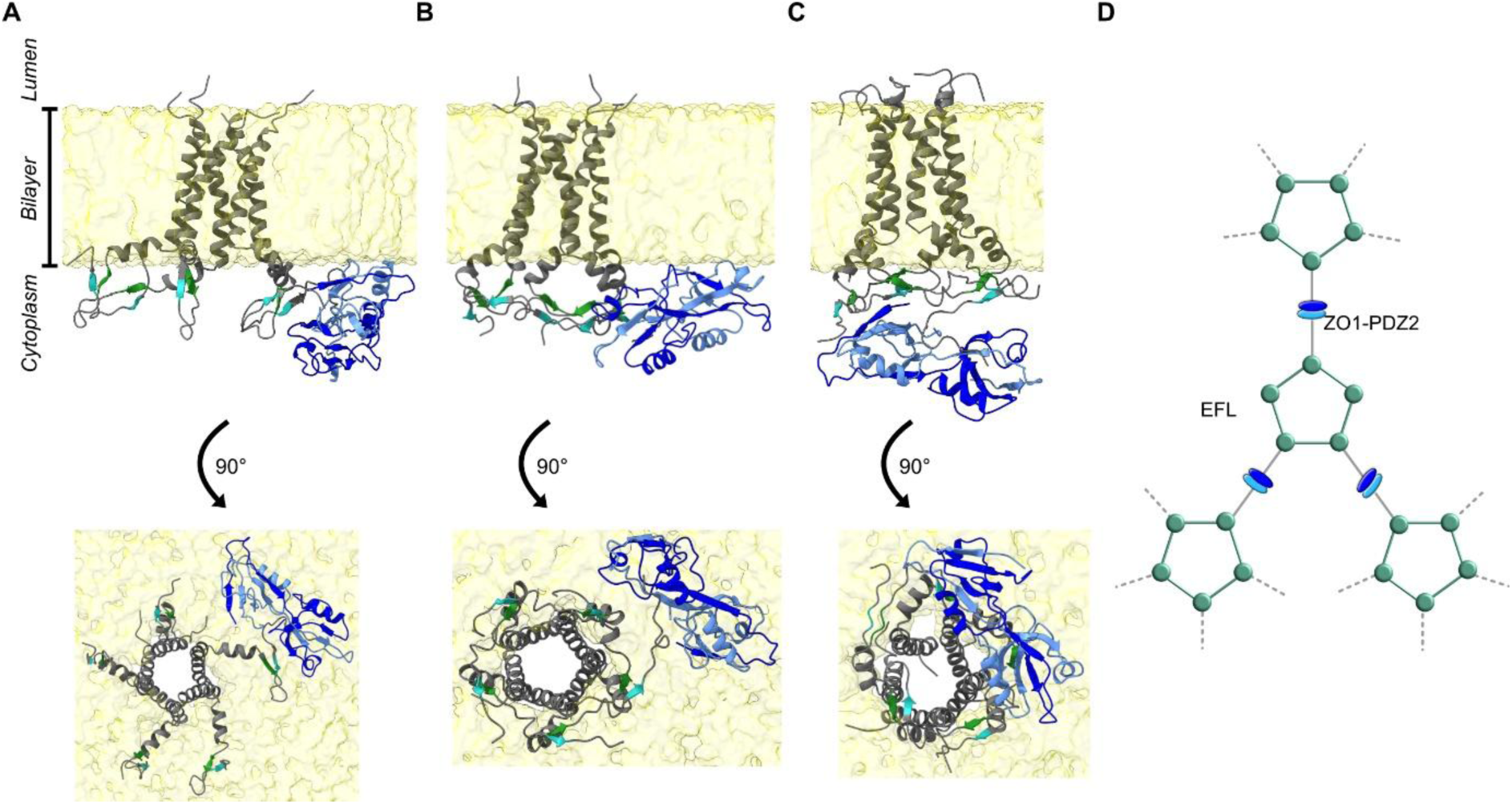
Structural models of pentameric EFL in complex with dimeric ZO1-PDZ2. **(A)** Structural model of interaction of E (grey) containing an antiparallel intra-monomer β-sheet is shown in complex with homodimeric ZO1-PDZ2 (blue and light blue) in a lipid bilayer (yellow). Residues 8 to 75 are displayed for the E monomer that interacts with one of the binding sites of the ZO1-PDZ2 homodimer while the other four monomers of E in the complex are displayed from residues 8 to 67. An orthogonal view is displayed below. The topology adopted by the protein in the ERGIC membrane is shown to the left.^4^ **(B)** Structural model with parallel inter-monomer β-strands is shown. The β1-strand of one monomer interacts with the β2-strand of a neighboring monomer in the clockwise direction when viewed as displayed below. All other features are similar to (A). **(C)** Similar to (B) but the β1-strand of the monomer interacts with the β2-strand of the other neighboring monomer in the anti-clockwise direction. **(D)** Cartoon model showing potential clustering of E pentamers due to cross-linking by ZO1-PDZ2.

The calculated structural models display possible arrangements of E within the complex and orientations of ZO1-PDZ2 with respect to the bilayer. The intra-monomer parallel β-sheet structural model places the α-helix B of the binding pocket of ZO1-PDZ2 facing upwards towards the bilayer surface, whereas in the two inter-monomer anti-parallel structural models, the orientation faces downwards away from the bilayer surface. Experimental evidence for the ZO1-PDZ2 orientation with respect to the lipid bilayer surface is required to choose the correct orientation. The amphipathic helix (residues 39 to 47) of E also shows various degrees of membrane insertion in the different models that would also need to be experimentally determined.

Based on the unsuccessful attempts at modelling ZO1-PDZ2 with both sites occupied by two E monomers from within the same pentamer, we speculate that ZO1-PDZ2 potentially acts as a cross-linker between oligomers (Figure 9D). This could increase clustering of pentamers of E and is supported by the clustering seen in TEM images (Supplementary figure 2D). The CODEX data (Supplementary figures 6A and 6B) showed that the addition of ZO1-PDZ2 to E increased the inter-pentamer distances compared to E without ZO1-PDZ2 at the same LPR of 20. This observation is consistent with increased steric occlusion between the pentamers due to the additional component. It can also be speculated that cross-linking by ZO1-PDZ2 promotes head-to-head interactions between E pentamers that results in less dense packing compared to head-to-tail interactions. This would imply greater phenylalanine distances and subsequently a slower CODEX decay.

Lastly, intracellular ZO1 has been shown to phase separate through the PDZ3-SH3-GuK (PSG) supra-domain^77^ that may increase the local concentration of E driven by cross-linking. Conversely, high expression levels of E could drive further phase separation by sequestering more ZO1. Thus, the two processes would act cooperatively to deplete ZO1 concentration at the TJ complex due to partitioning to the organellar membrane. This enhanced clustering propensity of E in organellar membranes may lead to changes in membrane curvature and contribute to the cellular remodeling seen in SARS-CoV-2 infected cells.^78^

## Conclusions

We have shown that full-length E from SARS-CoV-2 is a pentamer when reconstituted in an ERGIC-mimetic lipid bilayer. We introduced ZO1-PDZ2 into the above reconstitution and successfully co-assembled a host-viral protein complex *in vitro*. E maintained its pentameric oligomerization state within these rigid complexes and ZO1-PDZ2 was found to be immobilized on the bilayer surface due to its interaction with the PBM of E. NMR data on EFL also revealed that the protein contains β-sheet secondary structure in its CTD that appears to be pre-organized prior to the interaction with ZO1-PDZ2. We combined our findings with prior studies on E and ZO1-PDZ2 to generate a structural model of the protein complex. The models suggest that ZO1-PDZ2 acts as a cross-linker that promotes clustering of E pentamers.

## Methods

### Expression, purification and reconstitution

A synthetic DNA coding for EFL (Thermo Fischer) with all cysteine codons in the sequence mutated to alanine was cloned into a modified pET15 (Novagen) vector, resulting in the following sequence of EFL after thrombin cleavage, including 3 residues from the tag: GSHMYSFVSEETGTLIVNSVLLFLAFVVFLLVTLAILTALRLAAYAANIVNVSLVKPSFYVYSRVKNLNSSRVPDLLV. The F4Y/F56Y double mutant of this sequence was generated by PCR mutagenesis, along with a deletion of the histidine residue to shorten the scar from cloning. The fusion protein construct was expressed as inclusion bodies in the *E. coli* strain BL21(DE3) at 25 °C for 16 hours after induction with 0.5 mM Isopropyl-β-D-thiogalactopyranoside (IPTG). To produce ^15^N and uniformly ^13^C labelled protein, the expression was performed in minimal medium with ^15^N-NH_4_Cl as nitrogen source and ^13^C_6_-D-glucose as sole carbon source. For production of perdeuterated, uniformly ^13^C-labeled protein, the BL21(DE3) cells were stepwise adapted in minimal medium in D_2_O (Eurisotope), with deuterated (^13^C_6_-1,2,3,4,5,6,6-D_7_)-D-glucose (Sigma-Aldrich) as carbon source. To produce the ^15^N, ^13^C-γ-phenylalanine labeled, ^13^C-depleted F4Y/F56Y double mutant of EFL, the labelled amino acid was added 1 hour before induction to the medium with ^13^C-depleted D-glucose as carbon source. The protein was solubilized from the inclusion bodies with 20 mM Tris, pH 8.0, 300 mM NaCl, 5 mM imidazole, 1% (40 mM) Fos-Choline-16 (Anatrace), 10 % (w/v) glycerol, Complete-EDTAfree™ (Roche), 0.5 mM Phenylmethylsulfonylfluoride (PMSF) (Roth, Germany), Benzonase (Merck) and purified by metal affinity chromatography (Ni-NTA Agarose, Macherey Nagel, Germany). The fusion protein was incubated overnight at room temperature with thrombin protease in PBS at pH 7.4. The thrombin-cleaved protein was purified by reversed-phase HPLC^31^ and its integrity verified by electrospray mass spectrometry. The purified protein was lyophilized and refolded over night at room temperature at 200 µM final protein concentration in 50 mM NaH_2_PO_4_, pH 6.5, 25 mM NaCl, 40 mM Fos-Choline-16, 0.5 mM Pefabloc (Roth, Germany)).

A codon-optimized synthetic DNA fragment (Thermo Fisher), encoding the second PDZ domain of ZO-1 (amino acids T185 – E264) was cloned into modified expression vector pET16b with Tabac Etch Mosaic virus (TEV) cleavage site between the N-terminal Histidine tag and the target protein fragment. The expression construct was transformed into *E. coli* strain BL21(DE3). Protein expression was performed in Luria-Bertani (LB) medium for producing unlabeled protein, or in minimal medium supplemented with 1 g/L NH_4_Cl and 4 g/L D-glucose as the sole nitrogen and carbon sources, with ^15^N- and ^13^C labeling as required. For production of perdeuterated protein, the transformed BL21(DE3) cells were stepwise adapted in minimal medium in D_2_O and the minimal medium was supplemented with deuterated glucose ((^13^C_6_-1,2,3,4,5,6,6-D_7_)-D-glucose or (1,2,3,4,5,6,6-D_7_)-D-glucose, dependent on the labelling scheme). Protein expression was performed at 20 °C overnight after induction with 1 mM IPTG. The bacterial cell pellet from 1 L of expression culture was dissolved in 120 ml of buffer A (50 mM sodium phosphate, pH 7.0, 300 mM NaCl, 10 mM imidazole, 0.5 mM PMSF, with 1 tablet Complete EDTAfree^™^ (Roche)), supplemented with a spatula tip of each RNAse, DNAse and lysozyme. Cell lysis was performed by repeated sonication with stirring at 4 °C between the sonication steps. The protein was purified by metal affinity chromatography (Ni-NTA Agarose, Macherey-Nagel) and was incubated overnight at room temperature with TEV protease in phosphate buffer at pH 7.0. The TEV-cleaved protein was reloaded on Ni-NTA Agarose to remove the affinity tag and further purified by gel filtration on a Superdex 75-16-60 column (Cytiva). The protein was finally dialyzed against 50 mM sodium phosphate, pH 6.5, 25 mM NaCl, 0.5 mM PMSF. After adjusting the final protein concentration to about 23 mg/ml (2.4 mM) the protein was shock frozen in suitable aliquots and stored at -80 °C.

For complex formation of EFL with the ZO1-PDZ2 domain, the recombinant ZO1-PDZ2 fragment was added at 5:1 (mol:mol) ratio with respect to EFL. Subsequently, the complex was reconstituted into ERGIC-mimetic liposomes (1-palmitoyl-2-oleoyl-glycero-3-phosphocholine (POPC), 1-palmitoyl-2-oleoyl-sn-glycero-3-phosphoethanolamine (POPE), bovine phosphatidylinositol (PI), 1-palmitoyl-2-oleoyl-sn-glycero-3-phospho-L-serine (POPS), cholesterol at a molar ratio of 45:20:13:7:15)^34,42^ at an LPR of 20:1 (mol:mol). For this purpose, the protein-lipid suspension was dialyzed with two buffer exchanges at room temperature against NMR buffer (40 mM NaH_2_PO_4_, 40 mM glutamic acid, 40 mM arginine, 0.02% NaN_3_, pH 7.0). Finally, the detergent was removed with BioBeads (BioRad) by incubation overnight. The liposomes were collected by ultracentrifugation.

### MAS-NMR sample preparation

Samples for room temperature measurements were incubated in the NMR buffer containing an additional 100 mM copper EDTA to shorten the recycle delay through enhanced T_1_ relaxation. Each sample was centrifuged into a 1.3 mm ZrO_2_ rotor and sealed with FKM rubber plugs to prevent evaporation. The rotors were stored at 4 °C and remained stable over the course of 1 to 2 years.

For DNP samples, the NMR buffer was decanted and replaced by universal buffer (20 mM MES, 20 mM HEPES, 20 mM sodium acetate) containing 40 mM arginine, 40 mM glutamate at pH 7.2 since inorganic buffers have low pH stability at wide temperature ranges. Next, ^13^C-depleted glycerol (Cambridge Isotope Laboratories) was added to an approximate concentration of 60% glycerol (v/v) since the sample needs to be vitrified for efficient hyperpolarization. The sample was incubated in a sealed chamber with 30 mL 60% glycerol (v/v) to allow equilibration of concentration via vapor diffusion. Subsequently, ∼10 mM (460 μg) of a hybrid biradical^79^ was added to the sample as the source of DNP enhancement via the cross-effect.^80^. Finally, the sample was centrifuged into a 3.2 mM ZrO_2_ rotor and sealed with a silicone plug.

### Room temperature MAS-NMR experiments

ZO1-PDZ2 was assigned using (H)CANH, (H)CONH, (H)CA(CO)NH, (H)CO(CA)NH, (H)(CA)CB(CA)NH, (H)NCC-GODIST and (H)CC-GODIST sequences (Supplementary figure 4) that included homoTROP for the carbon to carbon polarization transfers.^44,81^ On isotopically-labelled EFL samples, (H)CANH, (H)CONH, (H)CONCA and 2D CC-RFDR spectra were initially recorded to assign the sharp peaks observed for EFL in complex with ZO1-PDZ2. (H)CACONH and (H)COCANH spectra were recorded to assign the broader resonances from EFL. The pulse sequences were coded using the SPEPS transfer element for carbon to nitrogen polarization transfers and homoSPEPS for the carbon-to-carbon polarization transfers (CA-CO transfers). Additionally, given that most of the resonances are overlapped in our sample, we traded a modest loss in resolution for additional sensitivity by curtailing the acquisition time in the indirect dimensions.

Unless otherwise indicated, MAS spectra were recorded on a Bruker Avance III HD narrow-bore spectrometer operating at 18.8 T (800 MHz) with a three-channel probe (^1^H,^13^C,^15^N) at a spinning frequency of 55 kHz in a 1.3 mm rotor, with the VT gas flow set to 1250 L/h. Measurements were also made using a Bruker Avance Neo spectrometer operating at 28.2 T (1200 MHz) using a three channel probe (^1^H,^13^C,^15^N) at 55 kHz MAS in a 1.3 mm rotor. The (H)NCC-GODIST and (H)(CA)CB(CA)NH spectra were recorded at a 14.1 T (600 MHz) Bruker Avance III HD spectrometer using a DVT600W2 BL1.3 mm HXY probe. ^31^P spectra were recorded on the same DVT600W2 BL1.3 mm HXY probe. Sample temperature in all cases was set to 245 K, which results in a sample temperature near room temperature.

All spectra involving SPEPS and homoSPEPS elements were pre-processed using Topspin au scripts for conversion from echo-antiecho to STATES-TPPI mode for the relevant dimensions. Spectra were acquired in blocks of approximately 24 hours and summed up after linear drift correction^82^.

### DNP-NMR experiments

DNP experiments were carried out on a 14.1 T (600 MHz) Bruker Avance III HD spectrometer operating at 14.1 T (600 MHz) with a 3.2 mm HCN probe at 94 K with a gas flow of 1200 L/h. Enhancement factors ranged from 20 to 30-fold, and buildup times were usually in the range of 4-6 s except for two samples: ^13^C-γ-phenylalanine, ^13^C-depleted, ^15^N-labelled EFL in complex with ^13^C-depleted-labelled ZO1-PDZ2 (1.3 s) and ^13^C, ^15^N-labelled EFL with ^15^N-labelled ZO1-PDZ2 (0.8 s) indicating relatively high biradical concentration (20-40 mM).

The CODEX and PDSD pulse sequences are shown in Supplementary figure 9 along with representative 1D and 2D spectra. The cryogenic temperature (100 K) that is used to efficiently transfer polarization from electrons to nuclei in DNP further aids in freezing out molecular motions such as ring flips that could otherwise interfere with the oligomer determination using CODEX.^83,84^ Additionally, unlabeled ^13^C sites present at 1.1% natural abundance can contribute unwanted additional dephasing of the CODEX curve. To minimize their contribution, we expressed the proteins from *E. coli* grown in a medium containing ^13^C-depleted (0.1%) glucose. Additionally, we used 2D ^13^C–^13^C correlation spectra as a reference instead of the conventional 1D CODEX reference spectra for further background correction, as shown before.^53^

### CODEX data analysis and simulation

CODEX simulations were carried out using the rate-matrix approach described by Hong and coworkers^46^. The elements of a rate matrix, *K̂*, comprises rate constants for magnetization exchange between two sites that are governed by the dipolar coupling strength and F(0). The rate matrix is propagated in time steps with the following equation:

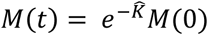

The dipolar coupling between different labelled sites were obtained from PDB file coordinates. A single global F(0) value was estimated for each sample by fitting the CODEX data to the simulations of using the four multimeric PDB models of E (Figure 2) using the reduced χ^2^ statistic as shown in Supplementary figures 6E, 6F, 6G and 6F and reported in Table S2.

The CODEX curve can decay less due to incomplete labelling. We therefore attempted to determine the labelling efficiency of the ^13^C-γ-phenylalanine in EFL using mass spectrometry. The TM segment of the chymotrypsin-digested protein did not elute due to the high affinity of the hydrophobic peptide to the C18 column. Since GB1 has been expressed with the same isotopic labelling, we quantified the labelling efficiency in GB1 instead, using mass spectrometry. GB1 showed 98% labelling efficiency (Supplementary figure 10). We believe that EFL would also show similar results. This labelling efficiency was also included in the simulation.^53^

### Data analysis of dipolar coupling measurement

The dipolar coupling is extracted by simulating the equations of motion and fitting to the data obtained with pMODERN recoupling (Supplementary figure 11)^59,85^. The measured dipolar coupling is converted to S by assuming a rigid NH bond length of 1.02 Å having a dipolar coupling strength of 11.48 kHz. S is subsequently converted to cone half-angle with the following formula:^61^

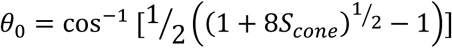

### Modeling parameters and protocol

Initial structural models of E and ZO1-PDZ2 in complex were generated using the NMR structure calculation software, CYANA. Three different types of restraints were used to inform various parts of the model: distance restraints, torsion angle restraints and residual dipolar coupling (RDC) restraints.

Distances between atoms were extracted from the TMD residues (12 to 38) of LMPG micellar ETR structure (pdb_00005X29) and converted to upper and lower distance restraints by including a ± 0.2 Å tolerance. Distances above 15Å were excluded. A similar procedure was applied for the E peptide bound ZO1-PDZ2 crystal structure (pdb_00009HM2) after using the ‘regularize’ module within CYANA to render the crystal structure compatible with CYANA. The regularization procedure moved the position of the peptide of E within the binding groove of ZO1-PDZ2 crystal, and therefore the distance restraints for the PBM and ZO1-PDZ2 interaction were extracted from the non-regularized structure. Approximately 22,000 distance restraints (N-N, CA-CA, CB-CB, CO-CO) were generated in this manner to fold the proteins *in silico*.

The amphipathic helix between L38 and N48, the two β-strands in the CTD and the single turn between S55 and V58 were modeled using torsion angle restraints. The assigned chemical shifts were used to predict the torsion angles using the software TALOS-N as mentioned before.^58^ For the residues in the predicted α-helix between L38 and N48, for which chemical shifts were not available, idealized φ (-60±10°) and ψ (-45±10°) angles for helices were used. α-helical segments were further restrained by hydrogen bonding restraints between the O atom of residue *i* and the H and N atoms of residue *i+4*. Similarly, parallel and anti-parallel β-sheets were generated by imposing distance restraints between residues in the two corresponding β-strands. Since the exact residues involved in hydrogen bonding between two β-strands is unknown, the strand register was chosen to maximize the number of hydrogen bonds in the resulting beta sheet.

The amphipathic helix was oriented approximately perpendicular to the TMD using RDC restraints within CYANA. Measured dipolar coupling values from oriented NMR samples^86^ can be used as RDC restraints in CYANA if the dipolar couplings are scaled down to Hz from kHz. The RDC values for the TMD were obtained from the dipolar coupling values reported by Zhang *et al.*^33^ and the RDC for amphipathic helix was set similar to the dipolar coupling values seen in the amphipathic helix of M2.^87^

The amphipathic helices between alternate monomers were also inhibited from close approach by setting a lower distance limit between residues that helped prevent the amphipathic helices from being positioned inside the pore of the pentamer. Similar repulsive distance bounds were also placed between residues in the TMD and the CTD to prevent the CTD from rising up towards the TMD since that would imply that the CTD is inserted into the lipid bilayer. ZO1-PDZ2 was also similarly prevented from fully inserting into the lipid bilayer by setting both RDC restraints and repulsive distance bounds from the TMD.

Simulated annealing under these restraints was performed in CYANA generating 1500 structures. A representative structure with a target function less than 10 was selected as a potential model of the E and ZO1-PDZ2 complex. The complex was then placed inside a lipid bilayer containing DMPC/DMPG (4:1 ratio) for visualization using the CHARMM-GUI web server^88^ after orienting the PDB using the OPM database web server.^89^

## Acknowledgements

We thank Kerstin Overkamp for HPLC purification and ESI-MS, Dr. Henning Urlaub and Dr. Olexander Dybkov at the Proteomics Facility and Dr. Dietmar Riedel at the Transmission Electron Microscopy facility at the MPI-NAT (Max-Planck Institute for Multidisciplinary Sciences) for their services. We further thank Dr. Vladimir Belov and Dr. Jan Seikowski from the Synthetic Chemistry Facility at the MPI-NAT for providing high-quality hybrid biradicals. We acknowledge funding from the Deutsche Forschungsgemeinschaft, Grants AN1316/1-2, AN1316/7-1, AN1316/8-1, and from the Max Planck Society.

