## Supplementary Information for "Bilayer-bound structure of SARS-CoV-2 Envelope protein in complex with a host PDZ domain"

### Materials and Methods

The samples used for NMR measurements and the corresponding pulse sequence details are listed below. The lipid composition is always ERGIC-lipids at LPR 20 unless mentioned otherwise.

Sample 1: $^{2}H$,$^{13}C$,$^{15}N$ labelled EFL (perdeuterated and 100% back exchanged) with $^{15}N$ labelled ZO1-PDZ2.

| **Spectrum** | **(H)CC-RFDR** | **(H)CANH** | **(H)CONH** | **(H)CANCO** |
|---|---|---|---|---|
| **Scans / points** | 320 | 8 | 16 | 64 |
| **Experimental time** | 3.5 | 22 hrs | 1.8 days | 4.8 days |
| **Field / MHz** | 1200 | 1200 | 1200 | 1200 |
| **Spinning frequency / kHz** | 55 | 55 | 55 | 55 |
| **Recycle delay / s** | 0.5 | 0.5 | 0.5 | 0.5 |
| **Sweep width (t1) / ppm** | 260 | 140 | 140 | 40 |
| **Maximum indirect evolution (t1) / ms** | 11.5 | 5.7 | 7.2 | 4.6 |
| **Sweep width (t2) / ppm** | — | 120 | 120 | 44 |
| **Maximum indirect evolution (t2) / ms** | — | 12.3 | 12.3 | 10.3 |
| Transfer I | H–C | H–CA | H–CO | H–CA |
| **$^{1}$H field / kHz** | 71.1 | 101.8 | 98.6 | 100.2 |
| **$^{13}$C field / kHz** | 70 | 42.2 | 42.2 | 42.8 |
| **Shape** | 90-100% linear ($^{1}$H) | 90-100% linear ($^{1}$H) | 90-100% linear ($^{1}$H) | 92-100% linear ($^{1}$H) |
| **Carrier ($^{13}$C) / ppm** | 120 | 60 | 180 | 53.7 |
| **Time / µs** | 2000 | 600 | 2100 | 800 |
| Transfer II | C–C | CA–N | CO–N | CA–N |
| **$^{13}$C field / kHz** | 83.33 | 10.7 | 10.7 | 22.6 |
| **$^{15}$N field / kHz** | — | 42.3 | 42.3 | 43.1 |
| **Shape** | RFDR | SPEPS[1] | SPEPS[1] | 63-94% tangent ($^{15}$N) |
| **Carrier ($^{13}$C) / ppm** | 120 | 60 | 180 | 53.7 |
| **Time / µs** | 1.16 | 2909 | 2909 | 7000 |
| Transfer III | — | N–$H^N$ | N–$H^N$ | N–CO |
| **$^{1}$H field / kHz** | — | 113.4 | 113.4 | — |
| **$^{13}$C field / kHz** | — | — | — | 53.7 |
| **$^{15}$N field / kHz** | — | 44.8 | 44.8 | 43.1 |
| **Shape** | — | 100-80% linear ($^{1}$H) | 100-80% linear ($^{1}$H) | 63-94% tangent ($^{15}$N) |
| **Carrier ($^{13}$C) / ppm** | — | — | — | 173.7 |
| **Time / µs** | — | 720 | 720 | 7000 |
| Transfer IV | — | — | — | — |
| **$^{1}$H field / kHz** | — | — | — | — |
| **X field / kHz** | — | — | — | — |

| | | | | |
|---|---|---|---|---|
| **Shape** | — | — | — | — |
| **Carrier ($^{13}$C) / ppm** | — | — | — | — |
| **Time / µs** | — | — | — | — |

| Spectrum | (H)CACONH | (H)COCANH | (H)COCACB | (H)NH-pMODERN[2] |
|---|---|---|---|---|
| **Scans per point** | 18 | 40 | 1248 | 64 |
| **Experimental time** | 11.6 days | 19.1 days | 21.2 days | 3.4 hours |
| **Field / MHz** | 800 | 800 | 800 | 800 |
| **Spinning frequency / kHz** | 55 | 55 | 55 | 55 |
| **Recycle delay / s** | 0.3 | 0.2 | 0.2 | 0.2 |
| **Sweep width (t1) / ppm** | 48 | 24 | 26.4 | 160 |
| **Maximum indirect evolution (t1) / ms** | 4.1 | 4.1 | 6.0 | 23.1 |
| **Sweep width (t2) / ppm** | 24 | 48 | 40 | — |
| **Maximum indirect evolution (t2) / ms** | 4.1 | 4.1 | 5.9 | — |
| **Sweep width (t3) / ppm** | 40 | 40 | — | — |
| **Maximum indirect evolution (t3) / ms** | 6.1 | 6.1 | — | — |
| Transfer I | H–CA | H–CO | H–CO | H–N |
| **$^1$H field / kHz** | 104.2 | 109.6 | 103.8 | 111.4 |
| **X field / kHz** | 39.7 | 51.6 | 38.0 | 41.4 |
| **Shape** | 80-100% linear($^1$H) | 80-100% linear($^1$H) | 80-100% linear($^1$H) | 80-100% linear($^1$H) |
| **Carrier ($^{13}$C) / ppm** | 48.7 | 168.0 | 178.7 | — |
| **Time / µs** | 3500 | 3500 | 3500 | 800 |
| Transfer II | CA–CO | CO–CA | CO–CA | N–$H^N$ |
| **$^1$H field / kHz** | — | — | — | 106.4 |
| **X field / kHz** | 52.5 | | 52.7 | 41.4 |
| **Shape** | homoSPEPS[3] | homoSPEPS[3] | homoSPEPS[3] | 100-80% linear($^1$H) |
| **Carrier ($^{13}$C) / ppm** | 108.7 | 108.0 | 113.7 | — |
| **Time / µs** | 1163 | 1163 | 1163 | 800 |
| Transfer III | CO–N | CA–N | CA–CB | — |
| **$^{13}$C field / kHz** | 11.5 | 11.8 | 68.8 | — |
| **$^{15}$N field / kHz** | 41.5 | 42.2 | — | — |
| **Shape** | SPEPS[1] | SPEPS[1] | homoSPEPS[3] | — |
| **Carrier ($^{13}$C) / ppm** | 168.7 | 48.0 | 62.7 | — |
| **Time / µs** | 2327 | 2909 | 1163 | — |
| Transfer IV | N–$H^N$ | N–$H^N$ | — | — |
| **$^1$H field / kHz** | 97.1 | 103.8 | — | — |
| **$^{15}$N field / kHz** | 41.1 | 41.7 | — | — |
| **Shape** | 100-80% linear ($^1$H) | 100-80% linear ($^1$H) | — | — |
| **Time / µs** | 650 | 500 | — | — |

The experimental parameters for (H)NH-pMODERN correspond to measurements for individual time points. The pMODERN mixing is incremented with a total constant time set to 1.018 ms.

Sample 2: $^{2}$H,$^{13}$C,$^{15}$N labelled EFL (perdeuterated and 100% back exchanged) with unlabelled ZO1-PDZ2.

| Spectrum | (H)NH |
|---|---|
| **Scans / points** | 16 |
| **Experimental time** | 4 hours |
| **Field / MHz** | 800 |
| **Spinning frequency / kHz** | 55 |
| **Recycle delay / s** | 1.2 |
| **Sweep width (t1) / ppm** | 200 |
| **Maximum indirect evolution (t1) / ms** | 19.9 |
| Transfer I | H–N |
| **$^{1}$H field / kHz** | 82.6 |
| **X field / kHz** | 36.7 |
| **Shape** | 80-100% linear ($^{1}$H) |
| **Time / µs** | 1300 |
| Transfer II | N–H$^{N}$ |
| **$^{1}$H field / kHz** | 75.1 |
| **$^{15}$N field / kHz** | 35.1 |
| **Shape** | 100-80% linear ($^{1}$H) |
| **Time / µs** | 700 |

Sample 3: $^{13}$C,$^{15}$N labelled EFL with $^{15}$N labelled ZO1-PDZ2.

| Spectrum | (H)CXCA(N)(CO)CX |
|---|---|
| **Scans / points** | 112 |
| **Experimental time** | 8.17 days |
| **Field / MHz** | 600 DNP |
| **Spinning frequency / kHz** | 12 |
| **Recycle delay / s** | 1 |
| **Enhancement** | 26X |
| **Sweep width (t1) / ppm** | 414 |
| **Maximum indirect evolution (t1) / ms** | 16.38 |
| **Sweep width (t2) / ppm** | 80 |
| **Maximum indirect evolution (t2) / ms** | 3.33 |
| **Sweep width (t3) / ppm** | 201 |
| **Maximum indirect evolution (t3) / ms** | 1.06 |
| Transfer I | H–CX |
| **$^{1}$H field / kHz** | 96.9 |
| **X field / kHz** | 69.4 |
| **Shape** | 90-100% linear ($^{1}$H) |
| **Carrier ($^{13}$C) / ppm** | 95 |
| **Time / µs** | 650 |
| Transfer II | CX–CA |
| **$^{13}$C field / kHz** | 0 (PDSD) |
| **$^{15}$N field / kHz** | — |
| **Shape** | — |

| | |
|---|---|
| **Carrier ($^{13}C$) / ppm** | 95 |
| **Time / µs** | 50,000 |
| Transfer III | CA–N |
| **$^{13}C$ field / kHz** | 46.8 |
| **$^{15}N$ field / kHz** | 30.5 |
| **Shape** | Square |
| **Carrier ($^{13}C$) / ppm** | 95 |
| **Time / µs** | 3000 |
| Transfer IV | N–CO |
| **$^{13}C$ field / kHz** | 47.7 |
| **$^{15}N$ field / kHz** | 30.5 |
| **Shape** | Square |
| **Carrier ($^{13}C$) / ppm** | 60 |
| **Time / µs** | 4200 |
| Transfer IV | CO–CX |
| **$^{13}C$ field / kHz** | 0 (PDSD) |
| **Carrier ($^{13}C$) / ppm** | 95 |
| **Time / µs** | 150,000 |

Sample 4: $^{2}H$,$^{13}C$,$^{15}N$ labelled ZO1-PDZ2 (perdeuterated and 100% back exchanged) with unlabelled EFL.

| Spectrum | (H)CC-GODIST[4] | (H)NCC | (H)NH-pMODERN[2] |
|---|---|---|---|
| **Scans / points** | 344 | 464 | 32 |
| **Experimental time** | 1.1 days | 9.1 days | 2.2 hours |
| **Field / MHz** | 800 | 600 | 800 |
| **Spinning frequency / kHz** | 55 | 55 | 55 |
| **Recycle delay / s** | 0.3 | 0.3 | 0.3 |
| **Sweep width (t1) / ppm** | 320 | 40 | 160 |
| **Maximum indirect evolution (t1) / ms** | 6.2 | 11.5 | 23.1 |
| **Sweep width (t2) / ppm** | — | 70 | — |
| **Maximum indirect evolution (t2) / ms** | — | 4.3 | — |
| Transfer I | H–C | H–N | H–N |
| **$^{1}H$ field / kHz** | 101.6 | 112.2 | 107.5 |
| **X field / kHz** | 38.4 | 42.8 | 41.4 |
| **Shape** | 80-100% linear ($^{1}H$) | 80-100% linear ($^{1}H$) | 80-100% linear ($^{1}H$) |
| **Carrier ($^{13}C$) / ppm** | 30 | — | — |
| **Time / µs** | 2500 | 700 | 800 |
| Transfer II | C–C | N–C | N-$H^N$ |
| **$^{1}H$ filed / kHz** | — | — | 104.2 |
| **$^{13}C$ field / kHz** | 27.3 | 12.9 | — |
| **$^{15}N$ field / kHz** | — | 40.9 | 41.4 |
| **Shape** | GODIST[4] | SPEPS[1] | 100-80% linear ($^{1}H$) |
| **Carrier ($^{13}C$) / ppm** | 30 | 54 | — |

| | | | |
|---|---|---|---|
| **Time / µs** | 3491 | 2909 | 800 |
| Transfer III | — | C–C | — |
| **$^{1}$H field / kHz** | — | | — |
| **X field / kHz** | — | 27.5 | — |
| **Shape** | — | GODIST[4] | — |
| **Carrier ($^{13}$C) / ppm** | — | -19 | — |
| **Time / µs** | — | 4655 | — |

The experimental parameters for (H)NH-pMODERN correspond to measurements for individual time points. The pMODERN mixing is incremented with a total constant time set to 1.018 ms. The rf-field strength corresponded to 138 kHz (~2.75*MAS).

| Spectrum | (H)CA(CO)NH | (H)CO(CA)NH | (H)(CA)CB(CA)NH |
|---|---|---|---|
| **Scans / points** | 64 | 96 | 160 |
| **Experimental time** | 2.2 days | 3.3 days | 9.3 days |
| **Field / MHz** | 800 | 800 | 600 |
| **Spinning frequency / kHz** | 55 | 55 | 55 |
| **Recycle delay / s** | 0.3 | 0.3 | 0.3 |
| **Sweep width (t1) / ppm** | 40 | 40 | 80 |
| **Maximum indirect evolution (t1) / ms** | 5.0 | 5.0 | 7.3 |
| **Sweep width (t2) / ppm** | 40 | 40 | 40 |
| **Maximum indirect evolution (t2) / ms** | 12.3 | 12.3 | 13.2 |
| Transfer I | H–CA | H–CO | H–CA |
| **$^{1}$H field / kHz** | 102.8 | 100.4 | 114.7 |
| **$^{13}$C field / kHz** | 38.4 | 38.4 | 42.3 |
| **Shape** | 80-100% linear ($^{1}$H) | 80-100% linear ($^{1}$H) | 80-100% linear ($^{1}$H) |
| **Carrier ($^{13}$C) / ppm** | 53.7 | 173.7 | 39.7 |
| **Time / µs** | 3000 | 3000 | 3000 |
| Transfer II | CA–CO | CO–CA | CA–CB |
| **$^{13}$C field / kHz** | 62.7 | 62.7 | — |
| **Shape** | homoTROP[5] | homoTROP[5] | — |
| **Carrier ($^{13}$C) / ppm** | 113.7 | 113.7 | — |
| **Time / µs** | 1800 | 1800 | — |
| Transfer III | CO–N | CA–N | CB–CA |
| **$^{13}$C field / kHz** | 12.8 | 12.8 | — |
| **$^{15}$N field / kHz** | 40.3 | 40.3 | — |
| **Shape** | SPEPS | SPEPS | — |
| **Carrier ($^{13}$C) / ppm** | 173.7 | 53.7 | — |
| **Time / µs** | 2909 | 2909 | — |
| Transfer IV | N–$H^{N}$ | N–$H^{N}$ | CA–N |
| **$^{1}$H field / kHz** | 105.1 | 105.1 | — |
| **$^{13}$C field / kHz** | — | — | 13 |
| **$^{15}$N field / kHz** | 43.1 | 43.1 | 40.9 |

| | | | |
|---|---|---|---|
| **Shape** | 100-80% linear ($^{1}$H) | 100-80% linear ($^{1}$H) | SPEPS |
| **Carrier ($^{13}$C) / ppm** | — | — | 53.7 |
| **Time / µs** | 500 | 500 | 2909 |
| Transfer V | — | — | N–H$^{N}$ |
| **$^{1}$H field / kHz** | — | — | 114 |
| **$^{15}$N field / kHz** | — | — | 42.8 |
| **Shape** | — | — | 100-80% linear ($^{1}$H) |
| **Time / µs** | — | — | 500 |

The (H)CANH and (H)CONH have the relevant transfer parameters as in the (H)CA(CO)NH and (H)CO(CA)NH. The out-and-back CA-CB-CA transfer in the (H)(CA)CB(CA)NH pulse sequence uses an INEPT element with 21.7 ms of $T_2$ delay.

### Supplementary Figures

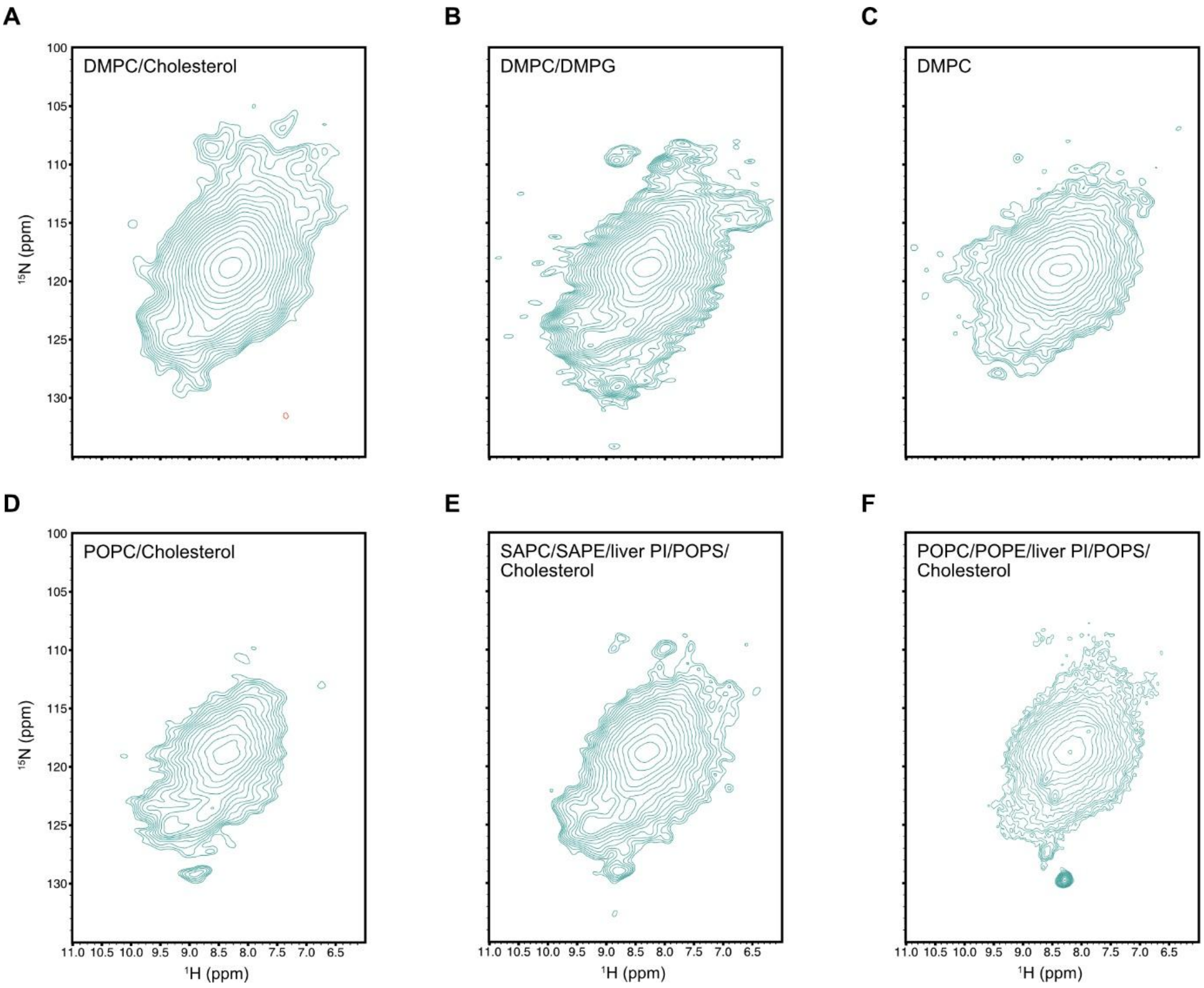


**Supplementary figure 1. $^{1}$H-$^{15}$N correlation spectra of EFL in various lipids.** EFL reconstituted with an LPR of 20 in **(A)** DMPC and cholesterol **(B)** DMPC and DMPG **(C)** DMPC **(D)** POPS and cholesterol **(E)** SAPC, SAPE, liver PI, POPS and cholesterol **(F)** POPC, POPE, liver PI, POPS and cholesterol. This five-lipid composition was chosen for all future experiments. All spectra were acquired at 800 MHz and 55 kHz MAS.

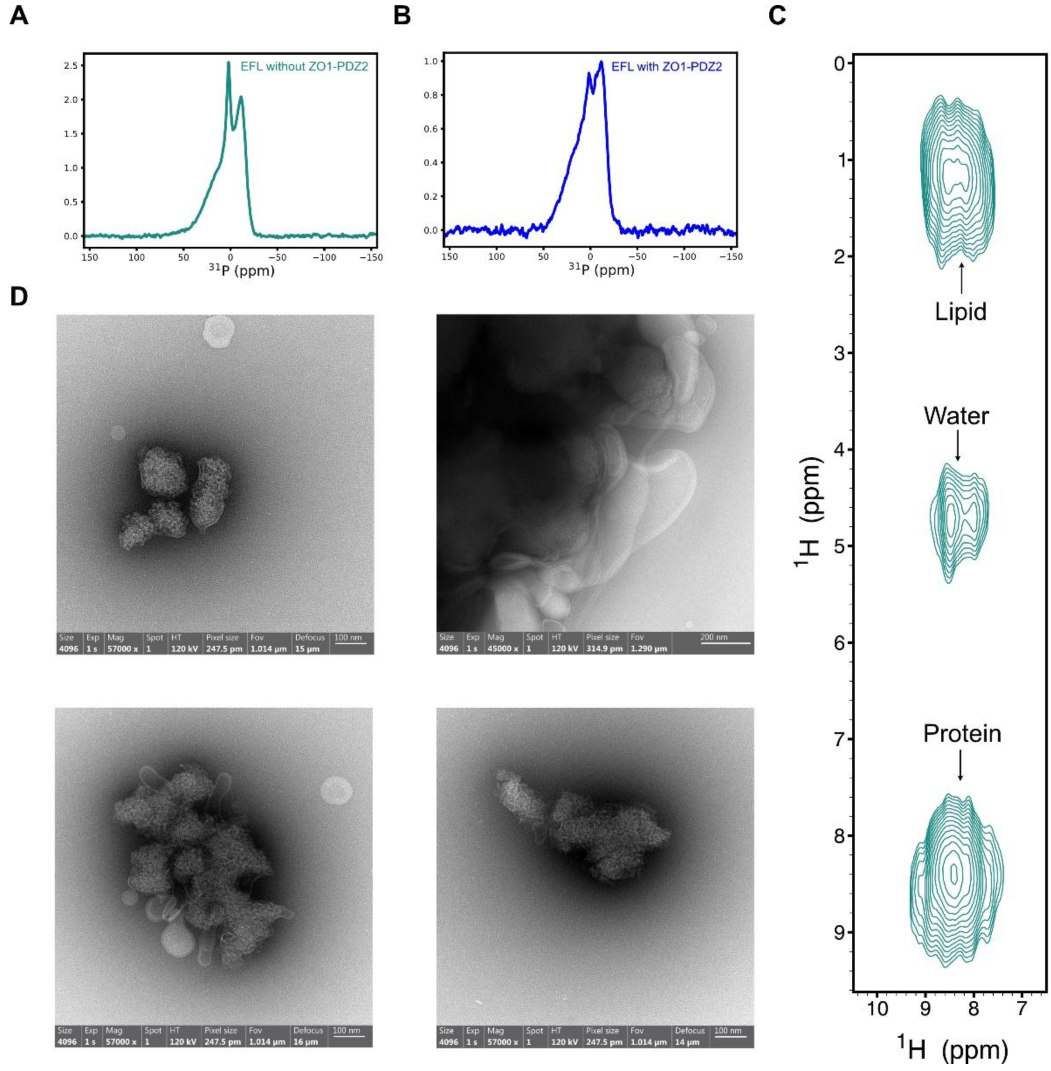


**Supplementary figure 2. EFL in liposomes (A)** $^{31}$P spectrum of lipid headgroups from liposomal preparation of EFL. The sharp peak corresponds to an isotropic peak from the phosphate buffer. **(B)** $^{31}$P spectrum of lipid headgroups from liposomal preparation of EFL in complex with ZO1-PDZ2. Both $^{31}$P spectra were acquired at 600 MHz without MAS. **(C)** Transmission electron microscopy images of EFL in complex with ZO1-PDZ2 in liposomes. **(D)** $^{1}$H-$^{1}$H correlation spectrum from an H(H)(CA)(N)H spectrum with $^{1}$H-$^{1}$H z-mixing showing overlapped cross-peaks between $^{1}$H/$^{13}$C/$^{15}$N-labelled EFL and alkyl chain $^{1}$H resonances from lipids confirming protein insertion during reconstitution acquired at 800 MHz at 55 kHz.

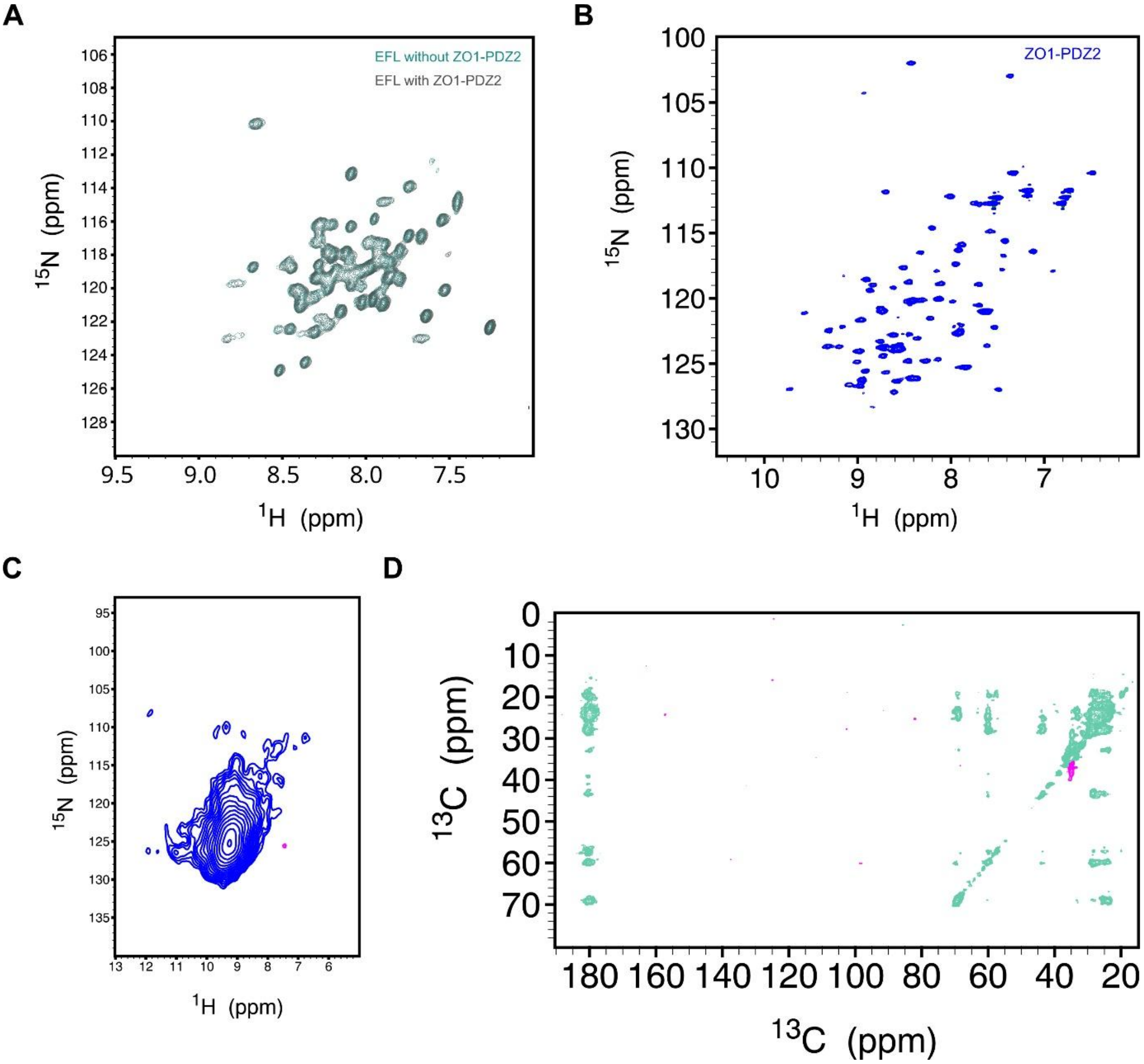


**Supplementary figure 3. ZO1-PDZ2 and EFL in different environments. (A)** Overlay of $^{1}H$-$^{15}N$ HSQC spectra of EFL solubilized in Fos-chol-16 with (grey) and without (green) ZO1-PDZ2. **(B)** $^{1}H$-$^{15}N$ HSQC spectrum of ZO1-PDZ2 in Fos-chol-16. **(C)** $^{1}H$-$^{15}N$ CP spectrum of $^{15}N$-labelled ZO1-PDZ2 reconstituted in ERGIC-lipids without EFL recorded at 800 MHz and 55 kHz MAS. **(D)** $^{13}C$,$^{15}N$-labelled EFL with ZO1-PDZ2 in 60%-glycerol and universal buffer at 600 MHz and 12.5 kHz MAS.

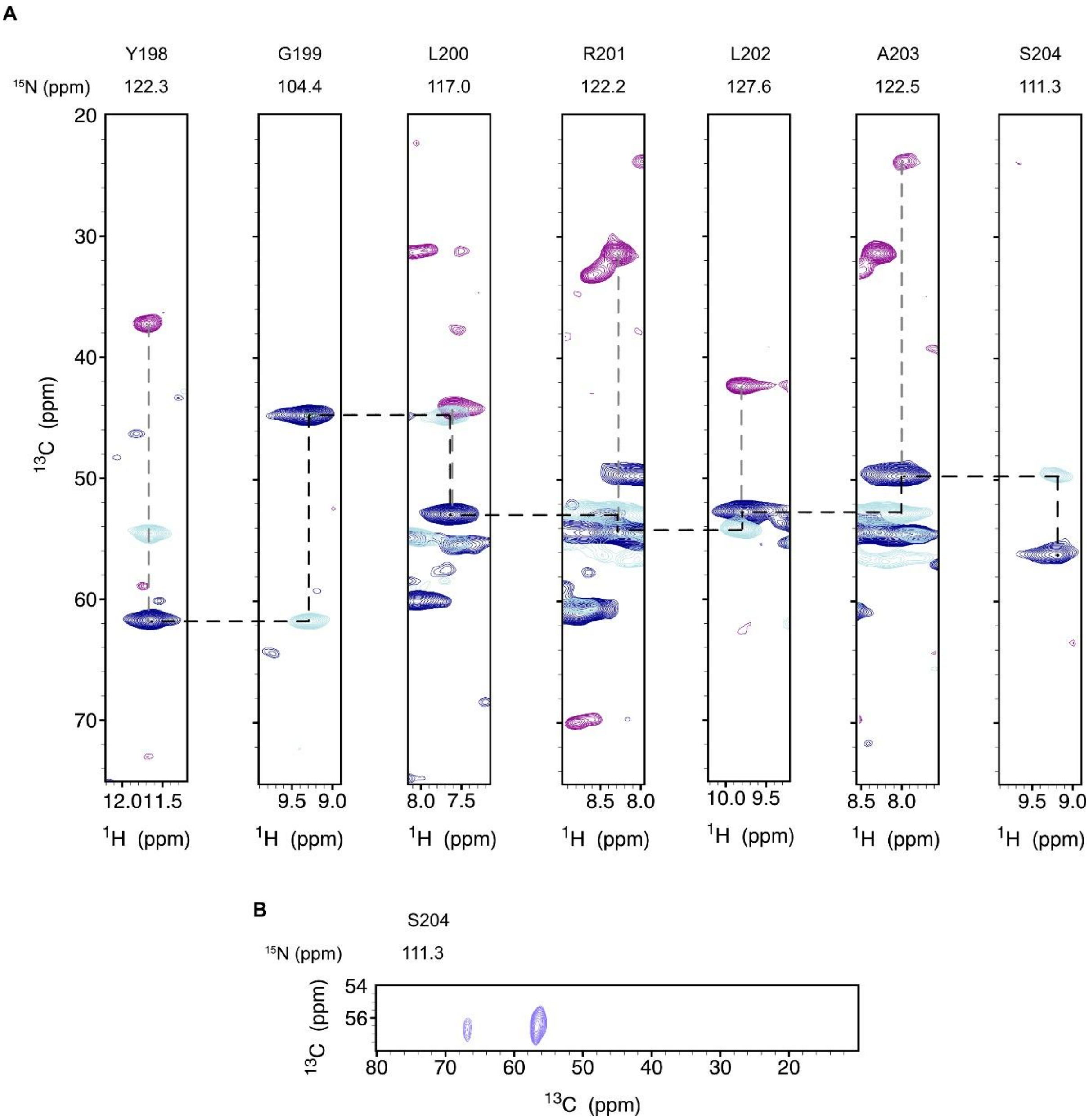


**Supplementary figure 4. Selected strips of assignment spectra of ZO1-PDZ2 (A)** Backbone walk with CANH (dark blue), CA(CO)NH (light blue) and CB(CA)NH (purple). **(B)** Missing side chain of S204 IN CB(CA)NH could be assigned from (H)NCC-GODIST spectra.

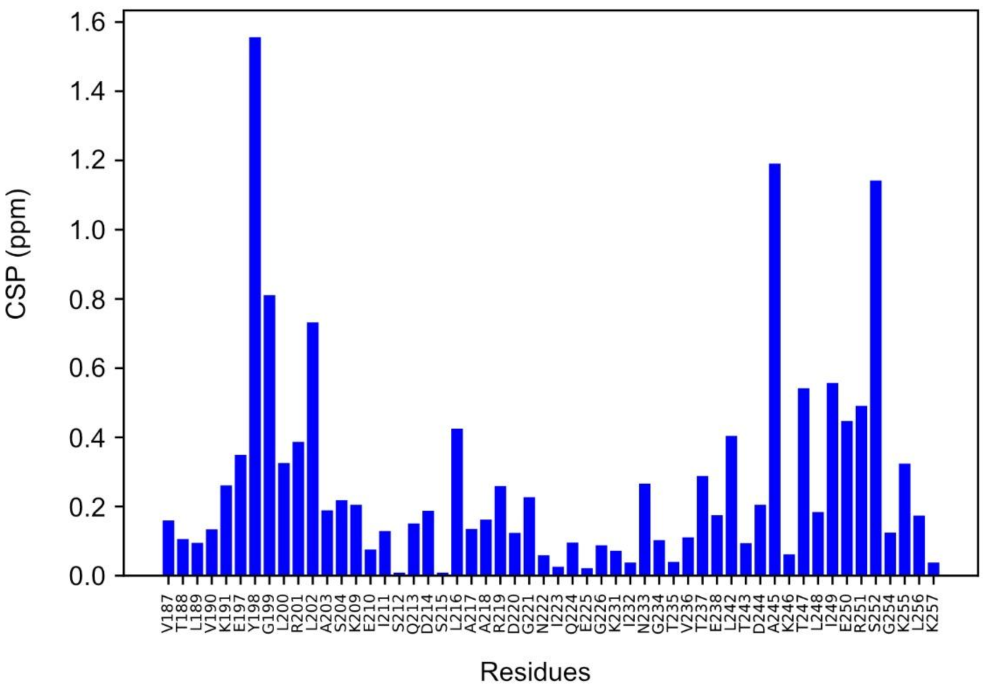


**Supplementary figure 5. Chemical shift perturbation (CSP) plot showing differences between solution NMR and MAS NMR assignments of ZO1-PDZ2.** The CSPs for all amino acids except glycine were calculated using the formula $\mathrm{CSP} = \sqrt{\Delta\delta_{\mathrm{H}}^2 + 0.14 \cdot \Delta\delta_{\mathrm{N}}^2}$, where $\Delta\delta_H$ and $\Delta\delta_N$ are the differences in proton and nitrogen chemical shifts, respectively, for a given peak. For glycine the formula was modified to $\sqrt{\Delta\delta_{\mathrm{H}}^2 + 0.2 \cdot \Delta\delta_{\mathrm{N}}^2}$.

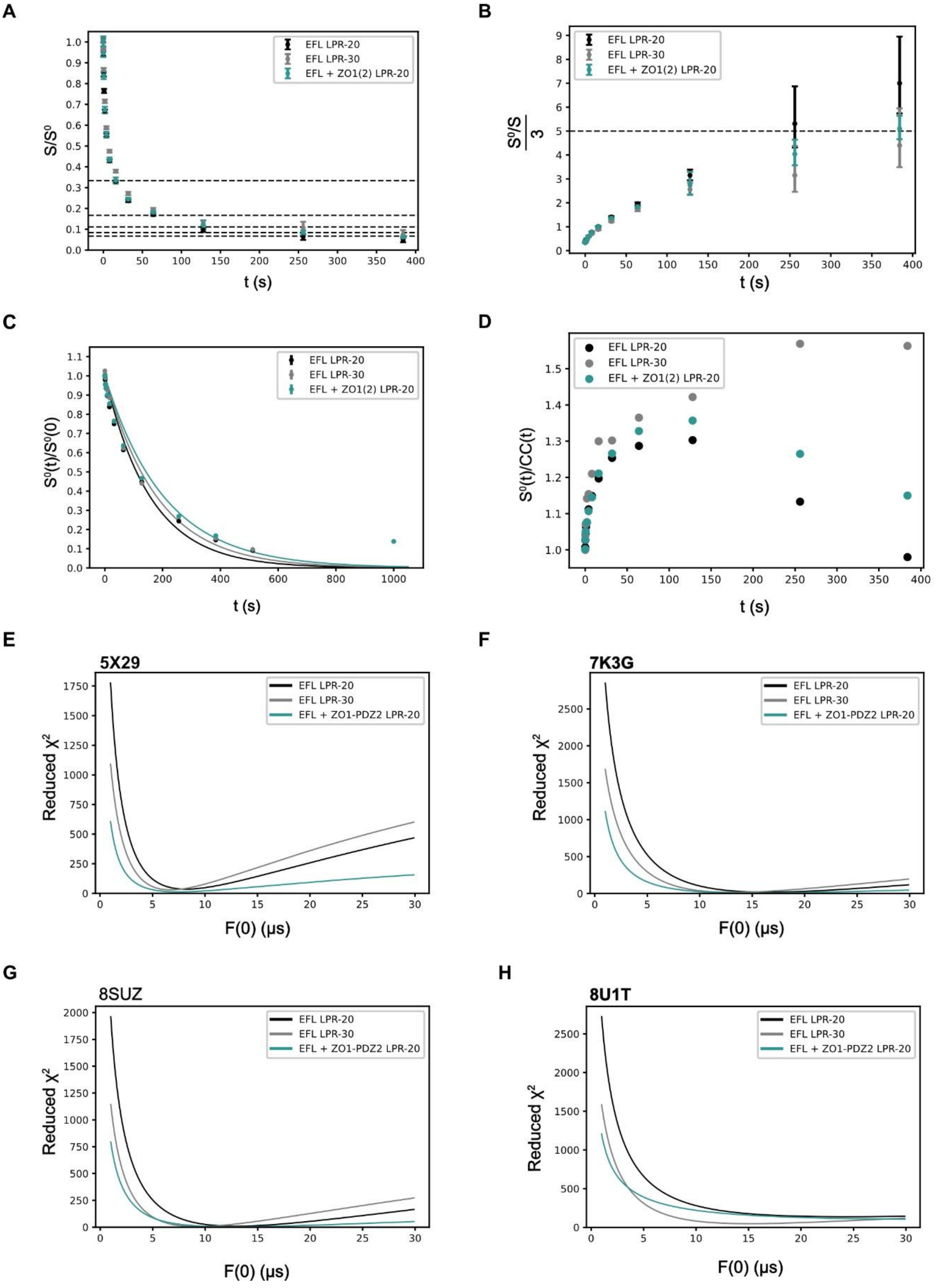


**Supplementary figure 6. CODEX data for EFL with and without ZO1-PDZ2. (A)** Corrected CODEX curves for EFL with ZO1-PDZ2 at LPR20 (cyan), and EFL without ZO1-PDZ2 at LPR 20 (black) and LPR 30 (grey) **(B)** The inverse plots of the CODEX dephasing curve normalized for three phenylalanine residues per monomer showing the corresponding oligomer numbers. **(C)** The normalized $S^0$ curves of the three samples overlap although their mono-exponential fits give different $T_1$ values of 181 s (EFL with ZO1-PDZ2 at LPR 20), 206 s (EFL without ZO1-PDZ2 at LPR 20) and 158 s (EFL without ZO1-PDZ2 at LPR 30). The poor fit for $T_1$ suggests multiexponential processes. **(D)** The correction plot shows the ratio of the CODEX reference curves in 1D ($S^0(t)$) and 2D (CC(t)). The correction factor plot indicates differences in spin diffusion rates to natural abundance background spins with a different isotropic shift compared to the labelled sites. **(E)** The reduced $\chi^2$ is plotted for different F(0) values fit to the CODEX data obtained on EFL with ZO1-PDZ2 at LPR 20 and EFL without ZO1-PDZ2 at LPR 20 and 30. Distances for the simulation were derived from the pdb_00005X29 structure. **(F)** Same as in (E) with distances derived from pdb_00007K3G. **(G)** Same as in (E) with distances derived from pdb_00008SUZ. **(H)** Same as in (E) with distances derived from pdb_00008U1T.

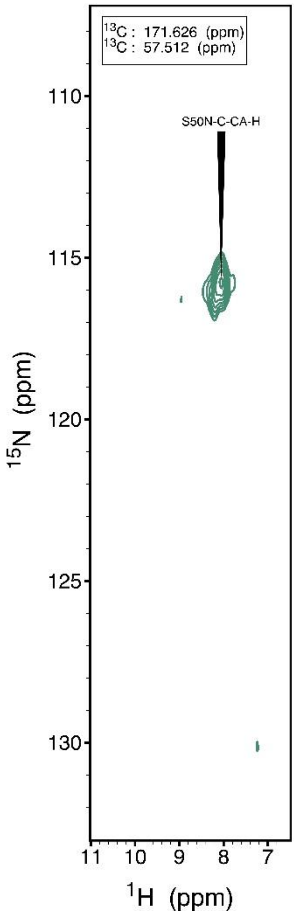


**Supplementary figure 7.** (H)COCANH spectrum of EFL in the absence of ZO1-PDZ2 showing the same S50 peak. The spectrum was acquired at 800 MHz at 55 kHz MAS.

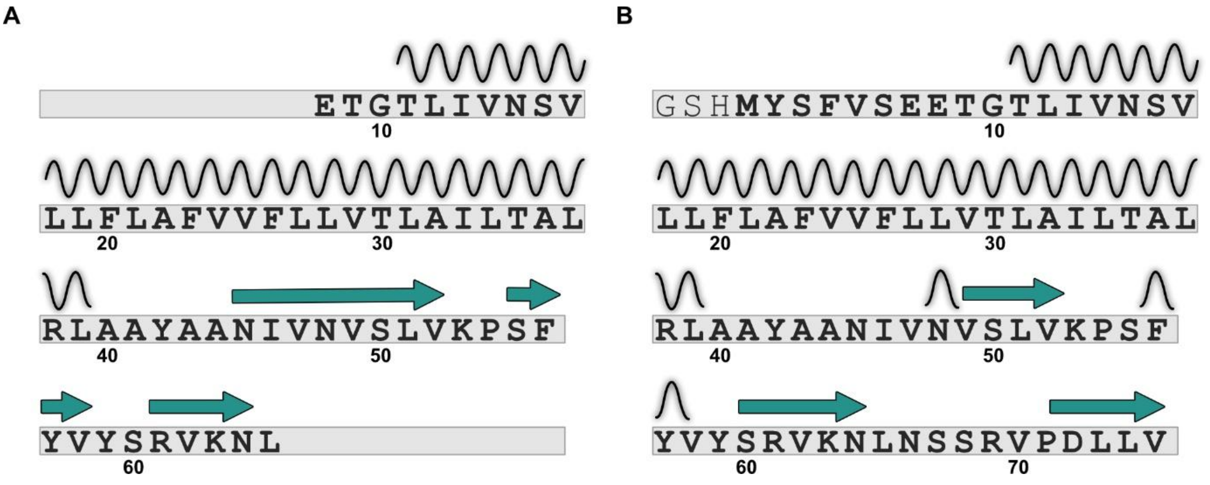


**Supplementary figure 8. Comparison of secondary structure of ETR and EFL. (A)** The sequence of $ETR_{8-65}$ is displayed with the corresponding secondary structures identified based on chemical shifts as reported by Dregni *et al.*[6] **(B)** The sequence of EFL is displayed with the corresponding secondary structures identified based on chemical shifts.

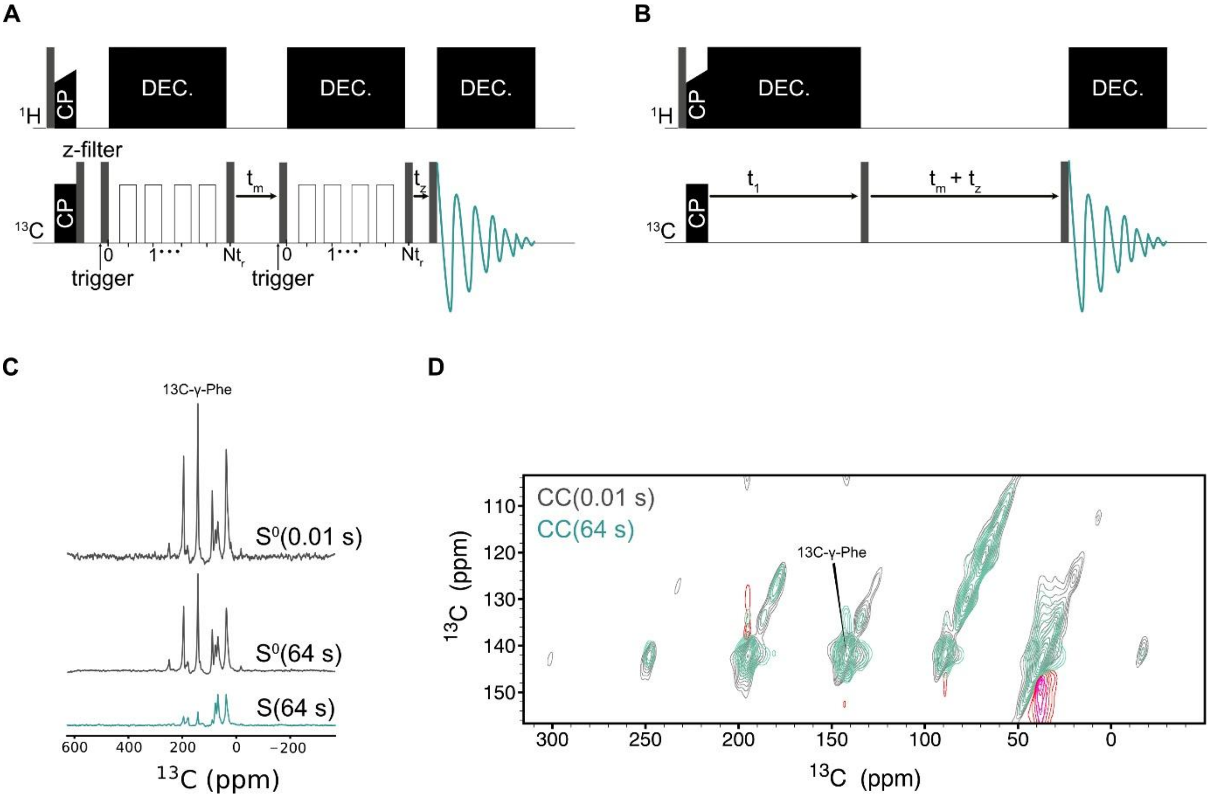


**Supplementary figure 9. CODEX and PDSD pulse sequence and spectra (A)** CODEX pulse sequence. $t_z$ was set to 10 ms. $t_m$ and $t_z$ were exchanged for the reference spectrum. **(B)** PDSD pulse sequence. **(C)** Representative CODEX spectrum of E with ZO1-PDZ2 showing the $S^0(0)$, $S^0(t)$ and $S(t)$ for 64 s time point. **(D)** Overlay of CC(0) and CC(t) $^{13}C$-$^{13}C$ PDSD spectra at 64 s which is used to correct the 1D CODEX spectrum. All spectra were acquired at 600 MHz at 8 kHz MAS.

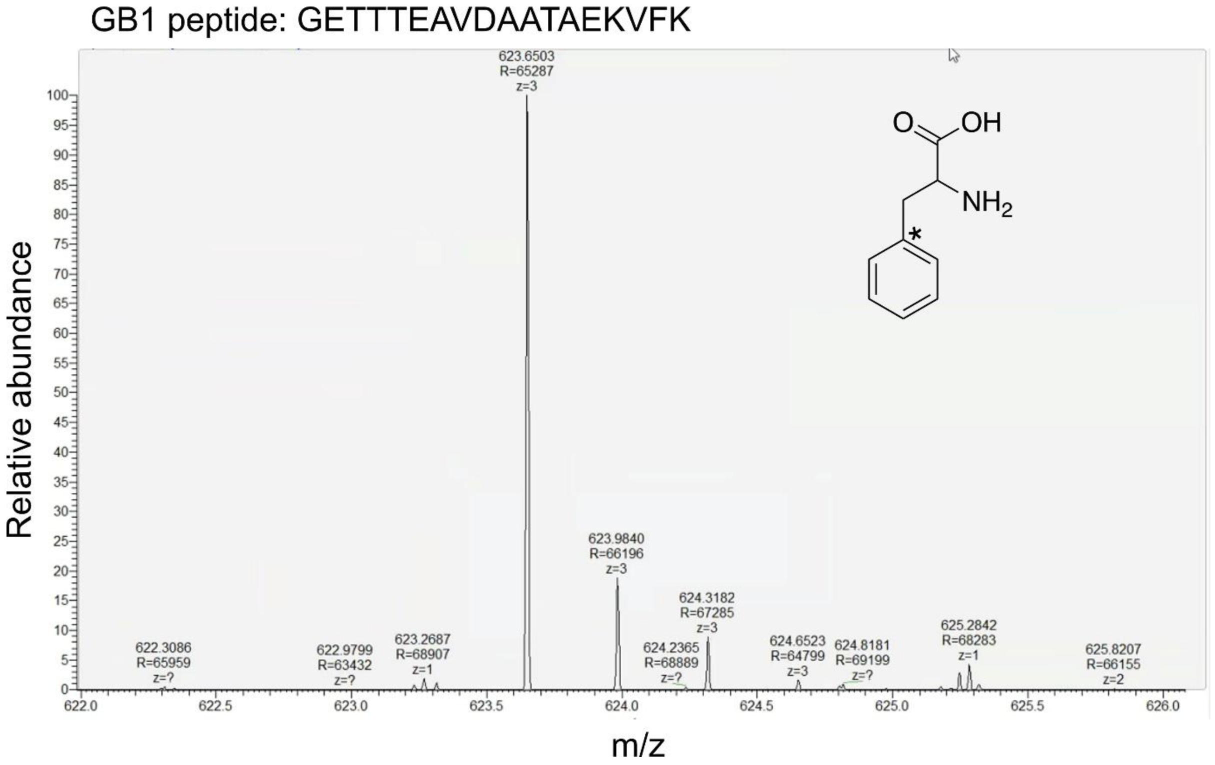


**Supplementary figure 10.** Isotopic-labelling efficiency of $^{13}$C-γ-phenylalanine was found to be 98% using GB1 expressed and purified under similar labelling media. The observed peptide in the mass spectrum after chymotrypsin-cleavage has the sequence shown above.

A

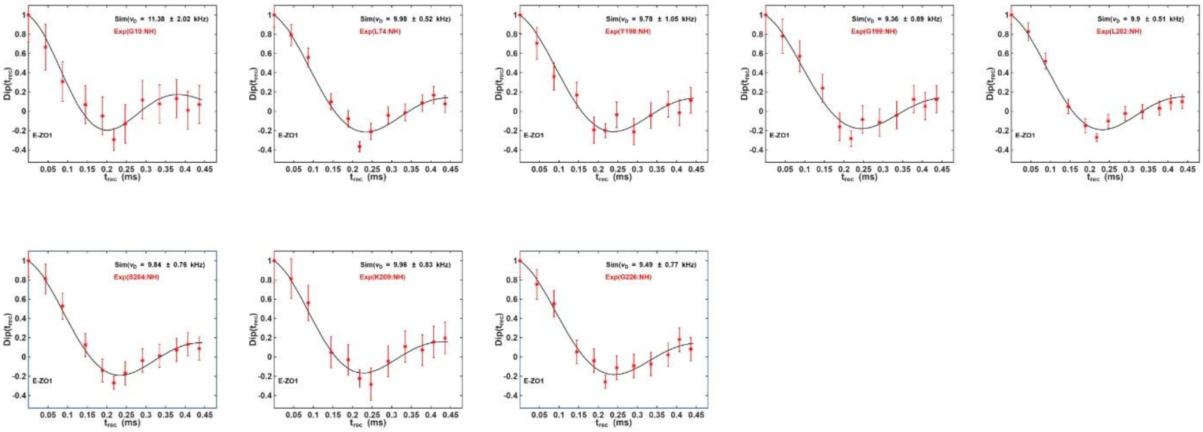
Dip($t_{rec}$)
$t_{rec}$ (ms)
Sim($\nu_D$ = 11.38 ± 2.02 kHz)
Exp(G10:NH)
Sim($\nu_D$ = 9.98 ± 0.52 kHz)
Exp(L74:NH)
Sim($\nu_D$ = 9.78 ± 1.05 kHz)
Exp(Y198:NH)
Sim($\nu_D$ = 9.36 ± 0.89 kHz)
Exp(G199:NH)
Sim($\nu_D$ = 9.9 ± 0.51 kHz)
Exp(L202:NH)
Sim($\nu_D$ = 9.84 ± 0.76 kHz)
Exp(S204:NH)
Sim($\nu_D$ = 9.96 ± 0.83 kHz)
Exp(K209:NH)
Sim($\nu_D$ = 9.49 ± 0.77 kHz)
Exp(G226:NH)
E-ZO1

B

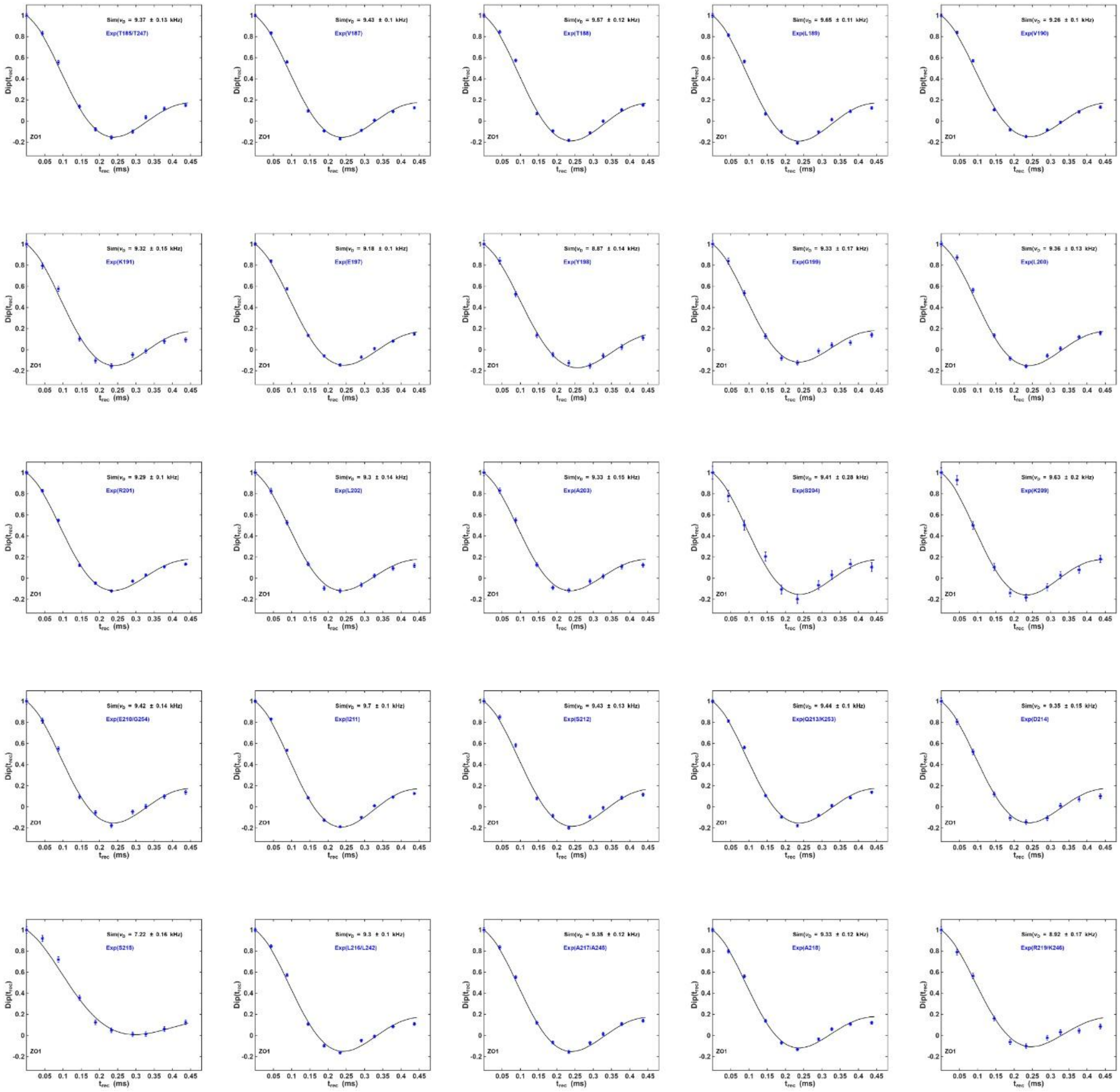
Dip($t_{rec}$)
$t_{rec}$ (ms)
Sim($\nu_D$ = 9.37 ± 0.13 kHz)
Exp(T185/T247)
Sim($\nu_D$ = 9.43 ± 0.1 kHz)
Exp(V187)
Sim($\nu_D$ = 9.57 ± 0.12 kHz)
Exp(T188)
Sim($\nu_D$ = 9.65 ± 0.11 kHz)
Exp(L189)
Sim($\nu_D$ = 9.26 ± 0.1 kHz)
Exp(V190)
Sim($\nu_D$ = 9.32 ± 0.15 kHz)
Exp(K191)
Sim($\nu_D$ = 9.18 ± 0.1 kHz)
Exp(E197)
Sim($\nu_D$ = 8.87 ± 0.14 kHz)
Exp(Y198)
Sim($\nu_D$ = 9.33 ± 0.17 kHz)
Exp(G199)
Sim($\nu_D$ = 9.36 ± 0.13 kHz)
Exp(L200)
Sim($\nu_D$ = 9.29 ± 0.1 kHz)
Exp(R201)
Sim($\nu_D$ = 9.3 ± 0.14 kHz)
Exp(L202)
Sim($\nu_D$ = 9.33 ± 0.15 kHz)
Exp(A203)
Sim($\nu_D$ = 9.41 ± 0.28 kHz)
Exp(S204)
Sim($\nu_D$ = 9.63 ± 0.2 kHz)
Exp(K209)
Sim($\nu_D$ = 9.42 ± 0.14 kHz)
Exp(E210/G254)
Sim($\nu_D$ = 9.7 ± 0.1 kHz)
Exp(I211)
Sim($\nu_D$ = 9.43 ± 0.13 kHz)
Exp(S212)
Sim($\nu_D$ = 9.44 ± 0.1 kHz)
Exp(Q213/K253)
Sim($\nu_D$ = 9.35 ± 0.15 kHz)
Exp(D214)
Sim($\nu_D$ = 7.22 ± 0.16 kHz)
Exp(S215)
Sim($\nu_D$ = 9.3 ± 0.1 kHz)
Exp(L216/L242)
Sim($\nu_D$ = 9.35 ± 0.12 kHz)
Exp(A217/A245)
Sim($\nu_D$ = 9.33 ± 0.12 kHz)
Exp(A218)
Sim($\nu_D$ = 8.92 ± 0.17 kHz)
Exp(R219/K246)
ZO1

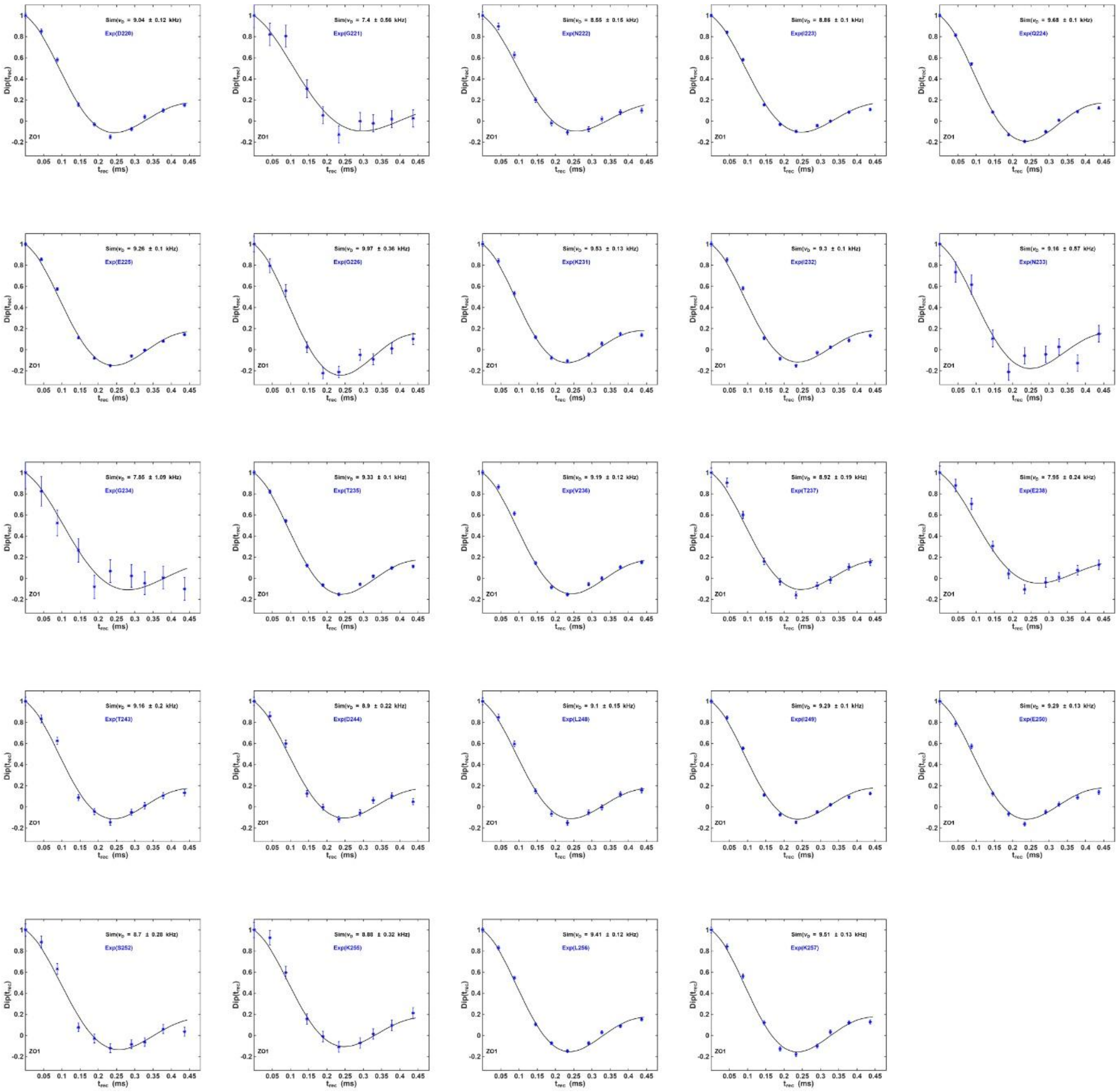

**Supplementary figure 11. $^{1}H$-$^{15}N$ single-bond dipolar coupling oscillation curves obtained with pMODERN recoupling. (A)** Residue specific dipolar oscillation curves are plotted as a function of the mixing time from a sample containing $^{2}H$,$^{13}C$,$^{15}N$-EFL and $^{15}N$-labelled ZO1-PDZ2. The first two plots with residues G10 and L74 correspond to peaks from EFL in complex with ZO1-PDZ2, while the remaining plots correspond to peaks from ZO1-PDZ2. **(B)** Residue specific dipolar oscillation curves obtained from $^{2}H$,$^{13}C$,$^{15}N$-labelled ZO1-PDZ2 without isotopic labelling on EFL. All spectra were acquired at 800 MHz at 55 kHz MAS.

**Supplementary Tables**

| | | |
|---|---|---|
| Gly 10 | N | 108.1 |
| Gly 10 | H | 8.4 |
| Gly 10 | CA | 47.4 |
| Gly 10 | CO | 174.7 |
| Asn 48 | N | 118.2 |
| Asn 48 | H | 8.2 |
| Asn 48 | CA | 61.1 |
| Asn 48 | CB | 40.6 |
| Asn 48 | CO | 176.4 |
| Val 49 | N | 118.8 |
| Val 49 | H | 8.2 |
| Val 49 | CA | 61.1 |
| Val 49 | CB | 38.2 |
| Val 49 | CO | 176.4 |
| Ser 50 | N | 115.9 |
| Ser 50 | H | 8.0 |
| Ser 50 | CA | 57.4 |
| Ser 50 | CB | 65.2 |
| Ser 50 | CO | 171.5 |
| Leu 51 | N | 121.4 |
| Leu 51 | H | 8.5 |
| Leu 51 | CA | 55.3 |
| Leu 51 | CB | 41.2 |
| Leu 51 | CO | 172.8 |
| Val 52 | N | 123.6 |
| Val 52 | H | 9.3 |
| Val 52 | CA | 60.3 |
| Val 52 | CB | 33.7 |
| Val 52 | CO | 174.1 |
| Tyr 57 | CA | 61.1 |
| Ser 60 | N | 124.2 |
| Ser 60 | H | 8.9 |
| Ser 60 | CA | 60.0 |
| Ser 60 | CB | 63.3 |
| Ser 60 | CO | 173.9 |
| Arg 61 | N | 125.3 |
| Arg 61 | H | 9.2 |
| Arg 61 | CA | 54.3 |
| Arg 61 | CB | 31.9 |
| Arg 61 | CO | 174.6 |
| Val 62 | N | 124.2 |
| Val 62 | H | 8.6 |

| | | |
|---|---|---|
| Val 62 | CA | 62.0 |
| Val 62 | CB | 32.4 |
| Val 62 | CO | 173.6 |
| Lys 63 | N | 125.4 |
| Lys 63 | H | 9.2 |
| Lys 63 | CA | 51.8 |
| Lys 63 | CB | 38.9 |
| Lys 63 | CO | 173.8 |
| Asn 64 | N | 125.1 |
| Asn 64 | H | 9.1 |
| Asn 64 | CA | 51.8 |
| Asn 64 | CB | 40.4 |
| Asn 64 | CO | 173.0 |
| Asp 72 | N | 116.7 |
| Asp 72 | H | 7.9 |
| Asp 72 | CA | 53.9 |
| Asp 72 | CO | 175.0 |
| Leu 73 | N | 121.7 |
| Leu 73 | H | 8.3 |
| Leu 73 | CA | 53.6 |
| Leu 73 | CB | 43.3 |
| Leu 73 | CO | 173.9 |
| Leu 74 | N | 129.6 |
| Leu 74 | H | 8.2 |
| Leu 74 | CA | 53.8 |
| Leu 74 | CB | 41.1 |
| Leu 74 | CO | 176.3 |
| Val 75 | N | 120.8 |
| Val 75 | H | 8.5 |
| Val 75 | CA | 60.5 |
| Val 75 | CB | 34.3 |
| Val 75 | COOH | 180.3 |

Supplementary Table 1: Assignments of E bound to ZO1(2)

| | EFL + ZO1-PDZ2 LPR 20 | | EFL LPR 20 | | EFL LPR 30 | |
|---|---|---|---|---|---|---|
| | **F(0)** | **red. $X^2$** | **F(0)** | **red. $X^2$** | **F(0)** | **red. $X^2$** |
| **pdb_00005X29** | 7.5 | 35.5 | 8.2 | 12.3 | 6.7 | 29.4 |
| **pdb_00007K3G** | 15.2 | 6.5 | 16.5 | 18.1 | 13.2 | 15.4 |
| **pdb_00008SUZ** | 12.8 | 2.9 | 12.9 | 9.0 | 9.2 | 4.8 |
| **pdb_00008U1T** | 29.9 | 104.7 | 24.4 | 137.0 | 15.1 | 47.3 |

Supplementary Table 2: Best-fit F(0) values and corresponding reduced $\chi^2$ values for the CODEX simulation with various PDB models. The F(0) values that fall within reported literature values are highlighted in grey.
